# Unified Mass Imaging Maps the Lipidome of Vertebrate Development

**DOI:** 10.1101/2024.08.20.608739

**Authors:** Halima Hannah Schede, Leila Haj Abdullah Alieh, Laurel Ann Rohde, Antonio Herrera, Anjalie Schlaeppi, Guillaume Valentin, Alireza Gargoori Motlagh, Albert Dominguez Mantes, Chloe Jollivet, Jonathan Paz-Montoya, Laura Capolupo, Irina Khven, Andrew C. Oates, Giovanni D’Angelo, Gioele La Manno

**Affiliations:** Institute of Bioengineering, School of Life Sciences, Swiss Federal Institute of Technology (EPFL), Lausanne, Switzerland; Brain Mind Institute, School of Life Sciences, Swiss Federal Institute of Technology (EPFL), Lausanne, Switzerland; Bioimaging and Optics Core Facility, School of Life Sciences, Swiss Federal Institute of Technology (EPFL), Lausanne, Switzerland; Center of PhenoGenomics, School of Life Sciences, Swiss Federal Institute of Technology (EPFL), Lausanne, Switzerland; Electrical Engineering Department, Sharif University of Technology, Tehran, Iran; Friedrich Miescher Institute for Biomedical Research, Basel, Switzerland

## Abstract

Embryo development entails the formation of anatomical structures with distinct biochemical compositions. Compared with the wealth of knowledge on gene regulation, our understanding of metabolic programs operating during embryogenesis is limited. Mass spectrometry imaging (MSI) has the potential to map the distribution of metabolites across embryo development. Here, we established an analytical framework for the joint analysis of large MSI datasets that allows for the construction of multi-dimensional metabolomic atlases. Employing this framework, we mapped the 4D distribution of over a hundred lipids at quasi-single-cell resolution in *Danio rerio* embryos. We discovered metabolic trajectories that unfold in concert with morphogenesis and revealed spatially organized biochemical coordination overlooked by bulk measurements. Interestingly, lipid mapping revealed unexpected distributions of sphingolipid and triglyceride species, suggesting their involvement in pattern establishment and organ development. Our approach empowers a new generation of whole-organism metabolomic atlases and enables the discovery of spatially organized metabolic circuits.

## Introduction

With the introduction and rise of single-cell and spatial omics methodologies, our capacity to describe cell-type composition and developmental trajectories has made a significant leap forward. Omics atlases continue to expand our understanding of the different factors influencing cell identity and positioning in both embryos and adult organisms [1–6].

Currently, compositional atlases rely on the ease of detection of nucleic acids and are built on the assumption that cell types and tissue structures are accurately described by their transcriptional profiles [5]. However, cell identity is not solely determined by gene expression, and organs are structured towards a metabolic division of labor [7, 8]. Therefore, mapping the detailed biochemical composition of tissues and organs is relevant, as it reveals an important axis of cell state heterogeneity and tissue structure.

In order to understand the biochemical tapestry of cells and tissues, a focus on lipids promises significant advances. Fundamental cellular processes such as energy metabolism, intracellular organization, transport and signaling rely on lipids [9–12]. Extracellularly, lipids act in paracrine and endocrine cell communication, contributing to the regulation of organism physiology [7, 13, 14].

Moreover, we have recently shown that single-cell lipid configurations drive cell states involved in tissue patterning [15]. During development, lipid levels display complex dynamics in vertebrates [16, 17]. Yet, despite their relevance, we have a limited understanding of how lipid compositional heterogeneity is organized in space and established at the organism level.

Mass spectrometry imaging (MSI) allows for the study of the spatial organization of lipids, as it measures the distribution of compounds at a micrometric scale, in parallel, and in an untargeted manner [18, 19]. An MSI acquisition is currently the measurement of choice to obtain an unsupervised biochemical description of a sample as a 2D image. However, only by combining multiple MSI acquisitions can one reveal a complete map that surveys lipids along organs and organisms (3D) over time (4D) and upon perturbations (**Figure 1a**).

**Figure 1:**
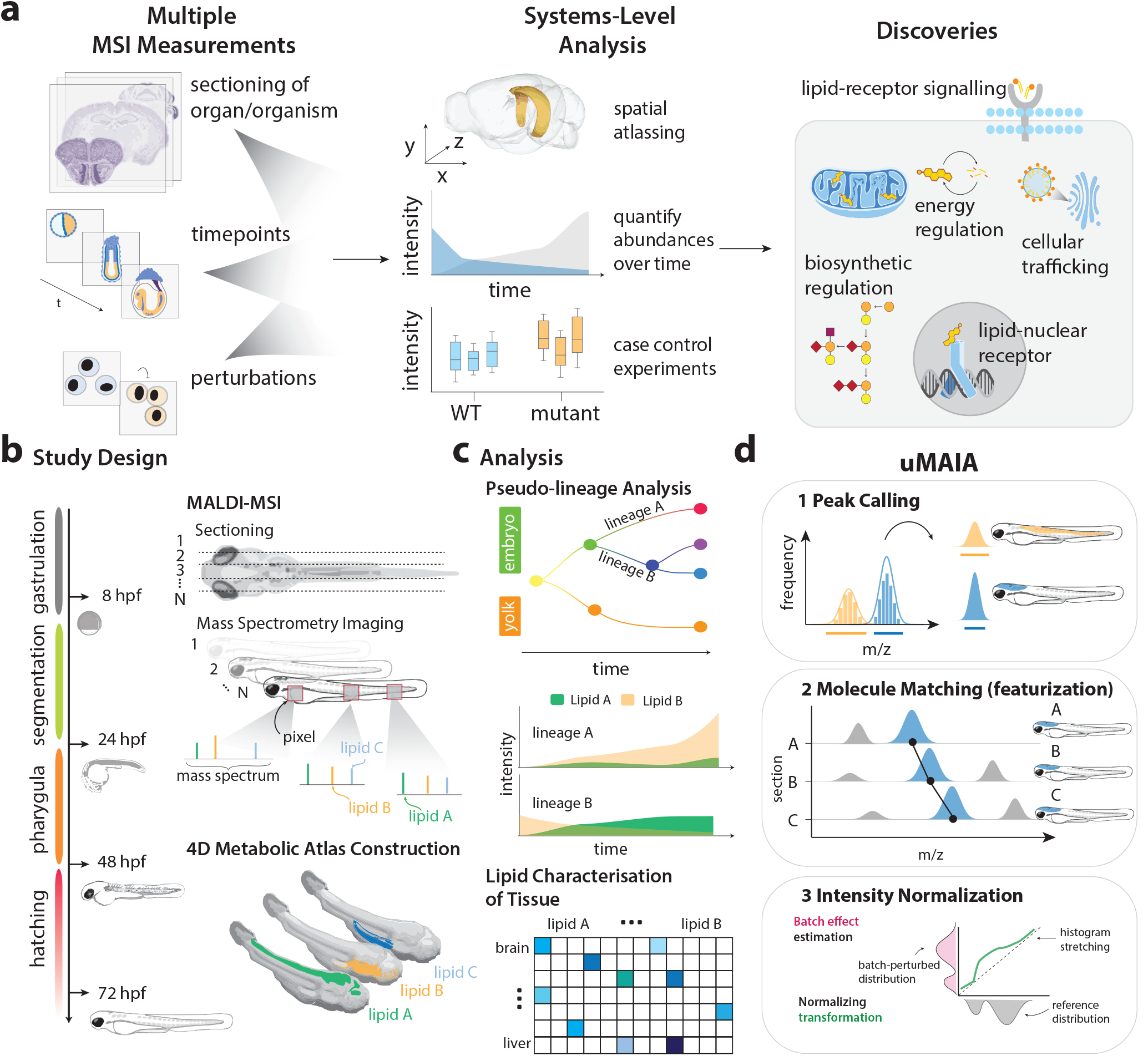
Workflow: from multiple MSI acquisitions to discovery using the uMAIA analysis framework. **(a)** General workflow enabled by uMAIA. From the left to the right: different MSI datasets consisting of collections of sections across organs, time points, or perturbations; aggregation and homogenization of the acquisitions to perform a system-level analysis; discovery of processes regulated region-specifically by lipids and metabolites. **(b)** Illustration of the experimental design. Zebrafish embryos at various developmental time points were prepared as a series of sagittal sections, and MALDI-MSI acquisitions were performed for each section independently. Construction of the 4D metabolic atlas. **(c)** Analysis of the 4D metabolic atlas, including regional and pseudolineage analysis and anatomical characterization of the 72 hpf zebrafish from the perspective of lipids. **(d)** Illustration of the 3 main steps of a typical processing with uMAIA: (1) Peak calling: images are extracted from single MSI acquisitions (2) Featurization by molecule matching: a unified feature space is obtained identifying, with an annotation free-approach, ions across multiple acquisitions and (3) Intensity normalization: experimental variability in intensity distribution is dampened preserving the signal.

We performed matrix-assisted laser desorption/ionization (MALDI)-MSI of entire vertebrate embryos at different stages of development to study the establishment of metabolically-defined anatomical zones (metabolic territories) (**Figures 1b, c**). We considered embryos from *Danio rerio* (more commonly known as zebrafish) that undergo substantial lipid remodeling during development [20, 21].

A few studies attempted to resolve the spatial localization of different lipid species in zebrafish. However, these analyses had limited spatial resolution or temporal scale, distinguishing just a few regions of the embryo (i.e., yolk, body, head, and tail) [22, 23] or focusing only on a small window of development [24]. A more detailed description of lipid spatiotemporal dynamics requires the analysis of high spatial-resolution MS images acquired throughout an entire organism and at different time points. However, large MSI datasets are challenging to integrate due to suboptimal image extraction, inappropriate molecule matching across samples, and intensity distortions (**Figure S1a**). Here, we devised a computational framework which we named the unified Mass Imaging Analyzer (uMAIA) to tackle these problems. uMAIA allows the generation of 4D lipid atlases of whole organisms starting from large collections of raw MSI acquisitions (**Figure 1d**).

With uMAIA we mapped a sizable fraction of the lipidome across zebrafish embryo anatomy and during development. We were able to study lipid spatiotemporal changes in the absence of external confounding factors, as zebrafish embryogenesis entails a closed metabolic system where energy sources and macromolecule building blocks are provided by yolk storages [8]. We traced the emergence of lipid territories that recapitulate embryo anatomical organization and found unexpected distributions of lipid classes that are relevant for tissue-specific functions. Altogether, our work demonstrates that the vertebrate developmental program involves the fine-grained remodeling of metabolism in a spatiotemporally regulated manner. It also highlights that lipid distribution is an accurate and information-rich descriptor of vertebrate anatomy, reinforcing the link between cell identity and lipid metabolism [25].

## Results

### The construction of a spatial lipidomic atlas is challenged by analytical and technical limitations

MSI data from multiple acquisitions are challenging to integrate due to technical variability in sample preparation and instrument performance. This manifests both as missing detections at the level of pixels or whole molecules and as non-biological deviations in signal intensity [26–28] (**Figure S1a**). These artifacts are common to MSI datasets, where the inconsistencies exist regardless of the specific MSI setting utilized and compound over acquisitions, hampering aggregated analyses of large MSI datasets (**Figure S1b-f**). *Ad hoc* measures, including the use of internal standards, have been devised to mitigate these effects, but they require prior knowledge of the sample composition, which is incompatible with untargeted approaches [28]. To overcome these limitations, we developed uMAIA. uMAIA enables analysis of large MSI datasets, as it accurately extracts images from spectra (peak calling), combines them in a shared feature space across acquisitions (featurization by peak matching), and subsequently intensity-normalizes them to minimize experimental fluctuations (normalization) (**Figure 1d**).

### Histogram-based adaptive peak calling allows for accurate mass image extraction

Image extraction from raw MSI data involves the identification of peaks generated by isobaric molecules across spectra (pixels). This process is complicated by instrument-driven fluctuations around the m/z value of a peak, known as mass shifts. Post-acquisition alignment methods have been developed to minimize these errors [29, 30]. However, calibration only partially corrects the problem [29–31] as we further evidenced in different analyses (**Figures 2a-c and S2a-e**).

**Figure 2:**
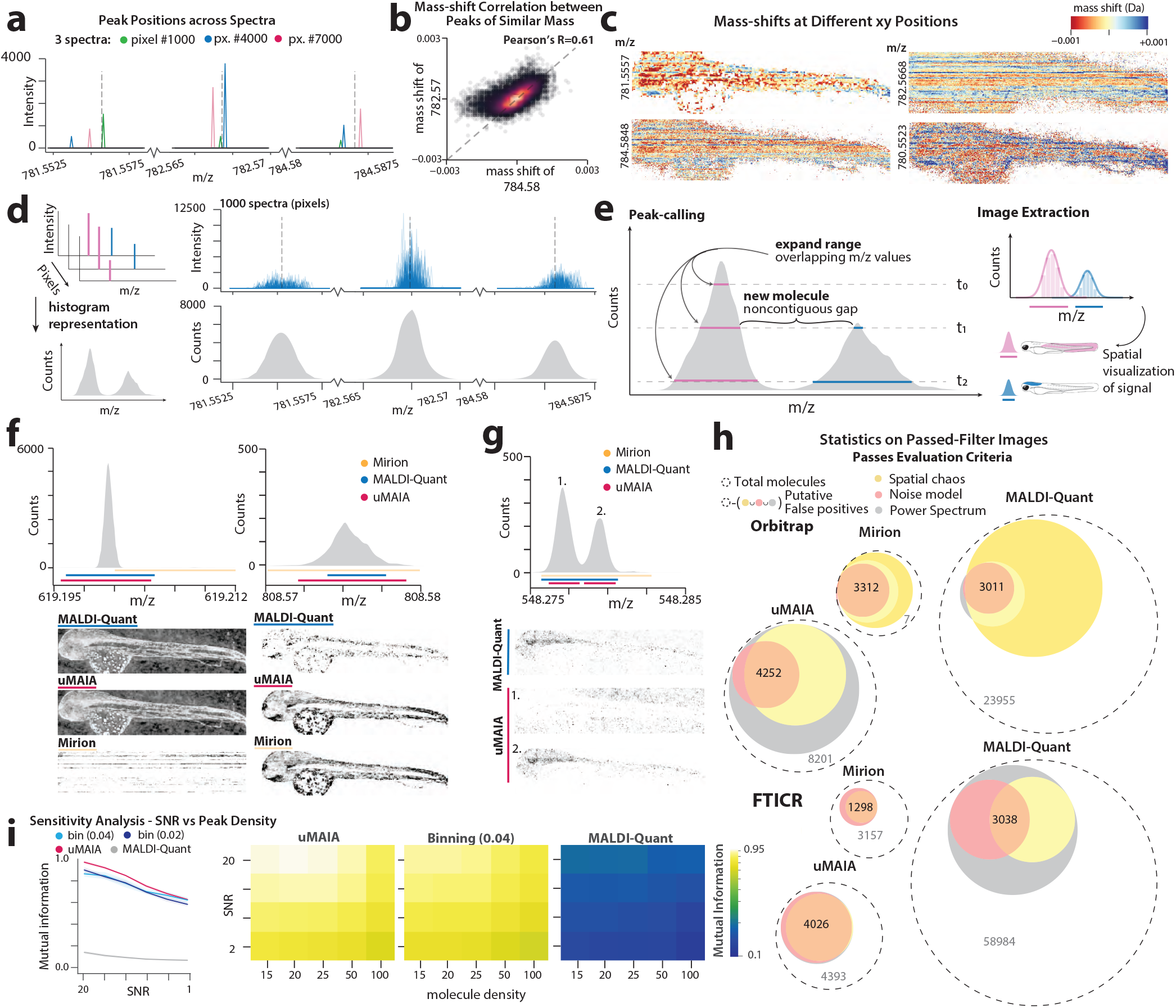
Performance comparison of uMAIA with other peak-calling algorithms on simulations and multiple datasets. **(a)** Peak positions (Da) and intensities for three representative molecules across 3 spectra. The subpanels are zoom-ins within m/z ranges that are in close proximity to each other and would, therefore, undergo a similar calibration adjustment with traditional methods. Vertical dotted lines correspond to the estimated m/z of each peak. Colors indicate the spectra (i.e., pixels) of origin. **(b)** Scatterplot between mass shifts of two molecules of similar mass (*∼* 2Da apart). Pearson’s R is indicated in the top right. **(c)** Heatmaps visualizing the intensity and direction of mass shifts in spatial context for the molecules shown in (b) along with two other molecules of similar mass. **(d)** Illustration of the first step of the approach used for peak calling, which reduces multiple spectra to a frequency (across pixels) histogram disregarding their intensity. On the right line, plots of 1000 spectra (i.e., pixels) are displayed stacked on the corresponding histograms for 3 representative cases. **(e)** Overview of uMAIA’s algorithm for peak calling by adaptive binning (left). The frequency histogram is used to identify bins. Intervals are iteratively expanded, iteratively decreasing a threshold (*t*_0_, *t*_1_, *t*_2_), as new intervals are considered. Peak intensity is integrated for each interval for each pixel to generate mass images (right). **(f)** Representative example of bins called using MALDI-Quant, Mirion, and uMAIA and the corresponding images generated. On top are histograms of the frequency of peak detections across the acquisition. Examples of three individual molecules are shown. Binning intervals are indicated by lines colored by the method used. On the bottom are the images extracted by the different methods for binning. **(g)** Histograms and images featuring a case of the different behavior among MALDI-Quant, Mirion and uMAIA in calling peaks: only uMAIA is able to discern two peaks, while the other methods merge them into one. Note the spatially coherent distribution of the two molecules, suggesting different compounds. **(h)** Sensitivity analysis on real data comparing uMAIA, MALDI-Quant, and binning for Orbitrap and FTICR MS acquisition. Venn diagrams depict the size of sets of molecules passing different assessment criteria used as a proxy for true positives (TP) in real data. Criteria used are the Spatial Chaos metric [36], assessment of the signal frequency (Power Spectrum; see Methods 3.1), and significance testing of a bivariate splines model (Noise Model; see Methods 3.1). The number within each Venn diagram is the number of high-confidence TP (e.g., as satisfied all three criteria). The gray number in the outer circle can be interpreted as a false positive estimate. **(i)** Left: line plot tracking the performance (mutual information score) of uMAIA, binning, and MALDI-Quant on simulated spectra. Average of 50 simulations; standard deviation over realizations represented as shaded area. Right: heatmaps display the average mutual information score for 50 simulation realizations as a function of both SNR and peak density.

Thus, to extract images from MSI data, m/z intervals must be considered that are assumed to encompass signals from isobaric molecules [32]. Commonly used methods retrieve intervals of a fixed size or whose extensions scale with m/z position [33, 34], but they do not account for the variability in peak-specific mass shifts which often results in artifacts. Adaptive approaches that attempt to encompass the mass shifts by retrieving intervals from intensity peaks have been proposed as a solution [35]. However, such approaches bias intervals to consider mainly peaks with high intensities even though lower-intensity peaks contain equally relevant spatial information. Therefore, we reasoned that counting the number of detection events at a position on the spectrum, disregarding intensity (i.e., the signal’s histogram representation), could better expose the distribution of mass shifts for individual peaks (**Figure 2d**). Based on this principle, we devised an approach where m/z intervals are initialized at the histogram maxima and then expanded until they either encounter other intervals or reach a background level of event counts (**Figure 2e**, see Methods 2.1). This procedure produces m/z intervals tailored to the spread of mass shifts characteristic of each individual peak.

We benchmarked our approach using data from different experiments and instruments against non-adaptive (Mirion [34]) and adaptive (MALDI-Quant [35]) methods. Inspection of images extracted by the different methods suggested that the uMAIA peak-caller effectively captured individual peaks (**Figure 2f** and **S2f-h**) and better resolved neighboring ones (**Figure 2g**). Indeed, uMAIA was remarkably more precise than Mirion in distinguishing quasi-isobaric compounds (2.3% vs. 42% images containing aggregated peaks) (**Figure S3a, b**). Conversely, MALDI-Quant tended to split intervals, generating multiple images that were similarly distributed and showed a checkerboard-like complementarity in space, suggesting the signal originated from a single ion (**Figure S3c**). Notably, peaks commonly detected by the different methods preserved peak centers and spatial distribution of the signal. Overall, uMAIA retrieved up to 55% more images of high quality (as defined in Methods 3.1) when compared to the second best method, depending on the MSI technology (**Figures 2h** and **S3d-f**). These findings were confirmed by studying the method’s behavior on simulated data where we controlled for different signal-to-noise ratios and peak density. uMAIA broadly outperformed existing methods, with a significantly greater signal recovery per peak than the next best method (Mutual Information Score uMAIA=0.98; binning=0.8; p-value«0.001) (**Figure 2i**).

### Network flow-based peak-matching places images in a unified and consistent feature space

To analyze datasets comprising more than one acquisition, peaks generated by the same molecule need to be identified (or ‘matched’) across different samples so as to featurize the data. The common way of accomplishing this task involves aligning detected spectra across samples using reference peaks, followed by naive binning (**Figure 3a, b**). This approach is relatively straightforward for a few samples and molecules but fails to generalize across the full repertoire of peaks in tens to hundreds of acquisitions, where spurious matchings become more likely. To obtain coherent matchings in large datasets, we conceived an unsupervised and alignment-independent approach that automatically links nearby peaks and groups them as the same molecule (**Figures 3a** and **S4a**). We consider the possible links between peaks and their mutual positioning to pose a Network Flow problem that can be efficiently optimized (see Methods 2.2 for a detailed description). This kind of formulation allows for setting constraints that prevent solutions that represent inconsistent scenarios, such as cases where a molecule has multiple matches originating from a single acquisition.

**Figure 3:**
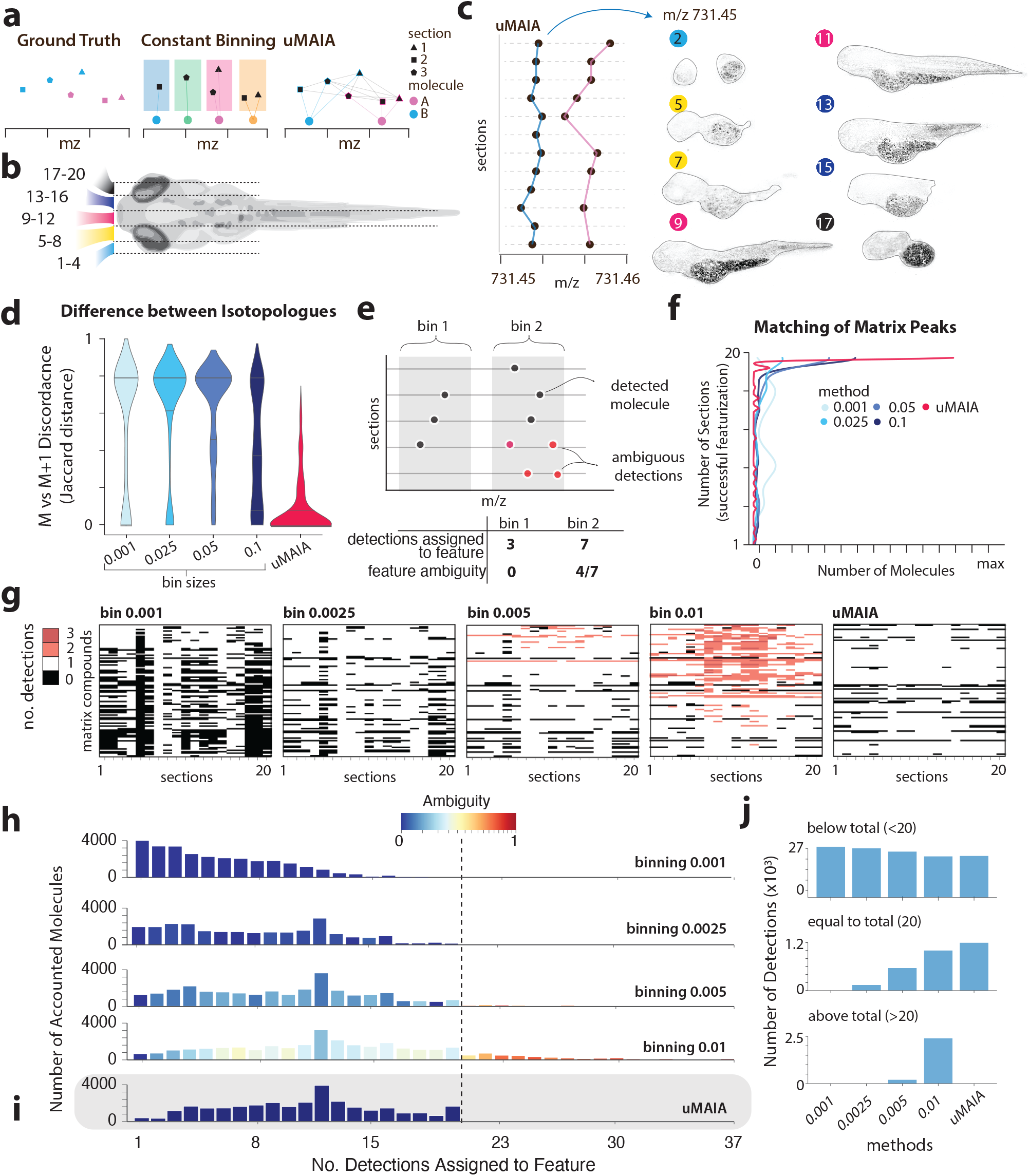
Assessment of the unified feature space by uMAIA’s peak matching and comparison with binning approaches. **(a)** Illustration of the peak-matching problem to place acquisitions into a coherent feature space. **(b)** Illustration of sagittal sectioning of a 72 hpf zebrafish to prepare MSI acquisitions. **(c)** Example of molecules matched across sections with detected m/z shown (left) and visualization of correspondent molecule, m/z=731.45 (right). **(d)** Violin plots reporting the distribution of Jaccard distances between isotopologue M+0 and M+1 presence across the acquisitions after featurization. Distributions are computed across all the pairs of sections for the 20 molecules with the highest signal intensity. **(e)** Illustration clarifying how feature ambiguity is defined and its score is calculated. **(f)** Density plots displaying the frequency of different peak-matching outcomes, the y-axis is the number of sections from the experiment in (b) where the featurization was successful. The histogram considers only the 50 most intense ions attributable to the MALDI matrix, so it is not confounded by biological variation. **(g)** Heatmaps visualizing the MALDI matrix ions (rows) identified for each section (column) for different binning sizes and uMAIA. Color-coded according to whether 0, 1, 2, or 3 peaks were identified within the bin for a given section. The same matrix compounds as in (f ) are displayed. **(h)** Barplots of the distribution of different peak-matching outcomes. The x-axis is the number of sections where the compound was identified across the entire dataset (20 sections). Bars are colored according to the average ambiguity within the range. Vertical dotted line indicates the total number of sections used (20). All compounds detected, irrespective of their intensity were used for this analysis. **(i)** As (h), corresponding to uMAIA-retrieved sets. **(j)** Barplots of the different peak-matching outcomes stratified by features: not detected in all sections (less than 20 sections), all sections (exactly 20 sections), or more than 20 sections for each method.

We tested our approach on simulated and real data and benchmarked it against binning procedures (see Methods 3.2). First, experiments on simulated data highlighted our method performed best for accuracy, recall, and precision and with outputs robust to the number of sections provided (**Figure S4b**). Next, we evaluated our approach on the zebrafish embryo acquisitions we had collected (**Figure 3b, c**). Due to the absence of a ground truth, we considered two cases where we could be confident about what to expect. In the first case, we took advantage of the signals from [M+0] and [M+1] naturally occurring isotopologues, which should be detected in the same acquisitions. Selecting the 20 most intense lipid peaks and identifying their [M+0] and [M+1] occurrences across the entire dataset indicated that uMAIA more consistently matched the detections of isotopologues in the same set of acquisitions compared to binning methods (**Figures 3d** and **S4c**). For the second case, we considered the 50 most intense MALDI matrix peaks (α-Cyano-4-hydroxycinnamic acid; CHCA) that are expected to be present across all acquisitions as MALDI matrix is applied to each sample. To evaluate the matching in this case, we introduced an ambiguity score to identify spurious matches by counting multiple detections from the same section that cannot, therefore, originate from the same molecule (**Figure 3e**). We quantified the occurrence of missing (dropouts) or multiple (ambiguous) matches using different binning sizes or uMAIA. We found that uMAIA maximizes the number of matched peaks while excluding ambiguous detections (**Figure 3f, g**).

We next asked to what extent peaks, irrespective of their source or intensity, could be matched across the entire dataset. When binning approaches were applied, the number of matched peaks and ambiguity scores scaled with bin sizes: smaller bins prioritized unambiguous detections at the expense of numbers of matched molecules, while increasing bin sizes retrieved larger groups but had higher ambiguity (**Figure 3h**). Our approach restricts groups to be unambiguous and retrieves a greater number of peaks across all acquisitions, corresponding to a 20% increase in detected signals (uMAIA=1200, binning=1000) than the largest bin size tested (0.01) (**Figure 3h-j**).

Thus, applying uMAIA is a robust way to extract peaks from multiple MSI acquisitions and consistently featurize them for downstream analyses. Importantly, uMAIA is annotation-free, thereby allowing one to carry out further analyses on a large portion of the data: in the case of our 72 hpf embryo MSI dataset, over 20,000 features were amenable to further analysis.

### uMAIA transforms intensity values to reduce technical variability in MSI data

Even when peaks are accurately called and matched, integrating MSI acquisitions remains challenging due to batch effects resulting from instrumental and experimental variability. This manifests as intensity distribution deformations [26]. While in the genomics field, many batch correction methods have been proposed [37–42], the data significantly differ from those acquired by MSI in terms of their signal distributions and type of batch effect.

We observed that when intensity values were logged, the MALDI-MSI data exhibited a bimodal distribution with low-(background) and high-(foreground) intensity modes (**Figure S5a**). Variable magnitudes of intensity distortions were observed across foreground modes in peaks detected in consecutive sections. This was the case even for matrix peaks that should have similar dynamic ranges across sections (**Figure 4a**). We hypothesized that the batch effects have factorizable components: a molecule-specific (*susceptibility*) and an acquisition-specific one (*sensitivity*). To investigate this aspect before developing a general estimation paradigm, we derive an *ad hoc* approximated empirical estimator of batch effect shifts. We exploited a property of our dataset, which is that true biological intensity should vary smoothly when moving through consecutive sections (see Methods 3.3). Then, we visualized the structure of the matrix of batch effect estimates (**Figure 4b**). The singular value decomposition confirmed the low rank of the matrix of empirically estimated batch effects and indicated that at least 31% of its values could be explained by its rank-one approximation, confirming the hypothesis of a strong factorization of the batch effect (**Figure S5b, c**). Based on this verified relationship, we devised a probabilistic model to perform a regularized estimate of these factors and remove non-biological distortion from the intensity distributions (**Figure 4c** and **S5d, e**, see Methods 2.3 for a detailed description).

**Figure 4:**
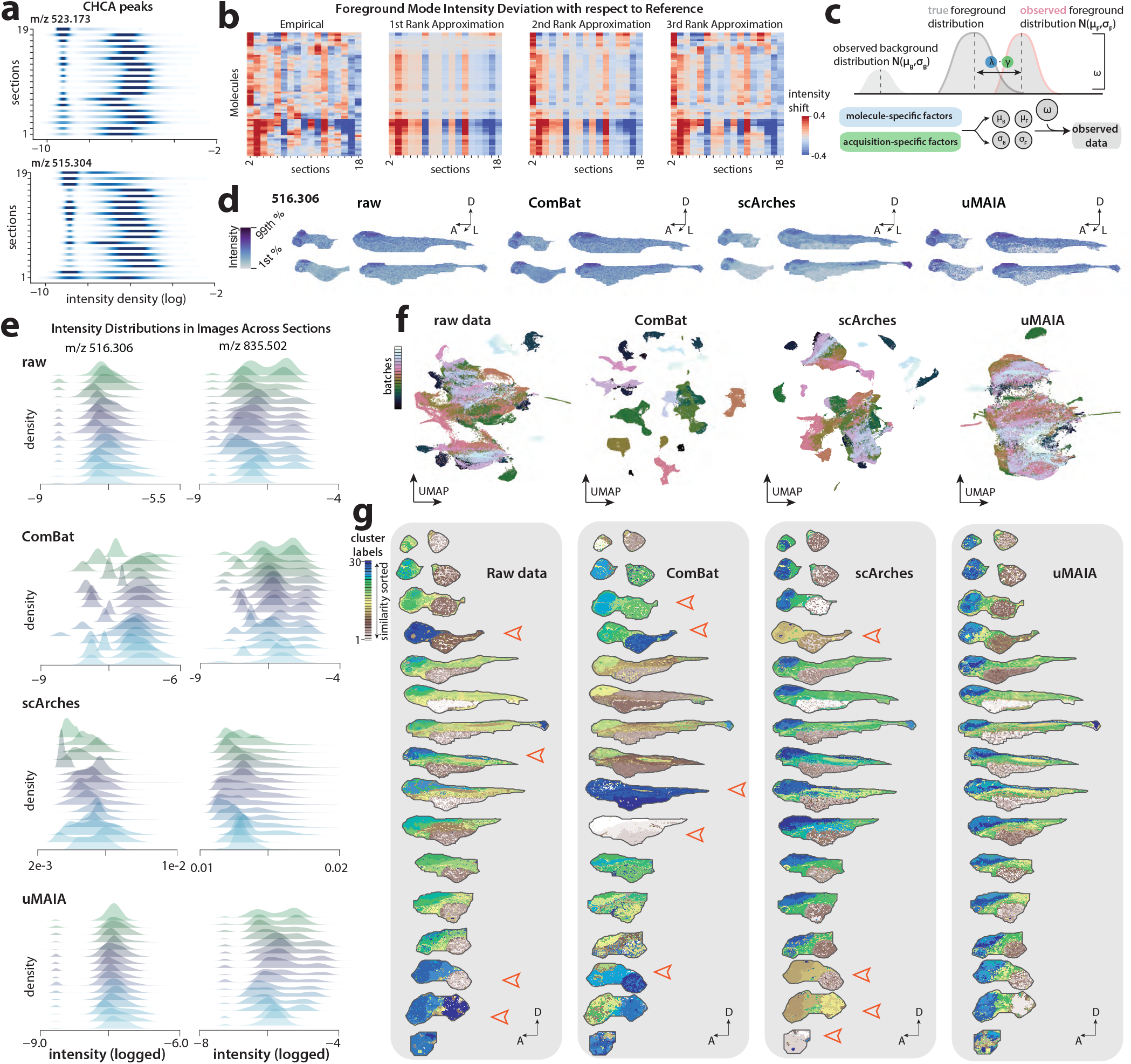
Characterization of technical variability in MSI data: intensity distortions and uMAIA’s normalization. **(a)** Intensity distribution densities of 2 CHCA (matrix) molecules across sections of the same embryo. The 72 hpf zebrafish atlas data was used. **(b)** Heatmap displaying the empirical estimates of foreground mode intensity shifts for 50 molecules (See Methods 3.3). Heatmaps on the right represent the 1st, 2nd and 3rd rank approximation of the matrix on the left. Approximations were obtained by the outer of the corresponding to groups of 1, 2, or 3 singular vectors, respectively. **(c)** Overview of the key modeling idea of factorizing decomposing distribution shifts. Signal distribution is approximated by a bimodal distribution (i.e., a mixture of two Gaussian) with a foreground and a background model. Observed foreground distribution parameters: center and standard deviation are subjected to a displacement with an offset, which we consider factorizable in two sources (molecule- and acquisition-specific: *λ* and *γ*, respectively). **(d)** Raw and normalized MSI images for representative molecules across landmark sections of the 72 hpf dataset. Colorbars are set to span between the 1st and 99th quantile of intensities across all sections. **(e)** Intensity distributions of molecules before and after normalization using different methods. **(f)** Low dimensional representation (UMAP embedding) of pixels for raw data, and the outputs of ComBat, scArches and uMAIA normalization. Pixels are color-coded by the section from which they originate. **(g)** Spatial visualization of discrete clusters after application of K-means algorithm on raw data and data processed with ComBat, scArches and uMAIA. Arrows indicate clear residual batch effects after clustering. The 72 hpf zebrafish atlas data was used.

First, we tested our approach on two simulated datasets where sections were perturbed with distortions that mimicked characteristics of the batch effect, as observed from real datasets (**Figure S5f**, see Methods 3.3). The simpler of the two was constructed to facilitate interpretation: overlapping 3D ellipsoids were generated and assigned different average signal intensities (**Figure S5g**). In addition, to challenge our model with more complex and realistic spatial distributions, we considered a second simulation based on patterns of expression extracted from the Allen Brain Atlas (ABA;100 genes, 16 coronal sections) [43] (**Figure S6**). We simulated batch effect intensity distortions for each section in both datasets, corrected the perturbed data with uMAIA, z-normalization, and ComBat, a popular location/scale method for batch-effect normalization [39], and evaluated the ability of the different algorithms to approximate ground-truth intensities (**Figures S5, S6c-h, S7**).

uMAIA performed well in all cases and it was both qualitatively and quantitatively superior to other methods in instances where the risk of stretching low-intensity values to signal was significant, displaying its ability to regularize the estimated parameters appropriately. This was indicated in the first dataset where uMAIA recovers the ordering of the modes of the raw data and reduces their variances, while ComBat and z-normalization intermixed the intensities of the distinct modes by assuming data to be Normally distributed (**Figure S5h-j**). The result was also reflected in the more complex ABA-based simulation: Root Mean Square Error (RMSE) scores between ground truth and normalized pixel values were lowest for uMAIA-corrected data (mean 89% improvement), followed by ComBat (mean 23% improvement), and worst yet highly gene-dependent for z-normalization (mean 17% decline) (**Figure S6c**). Additionally, performance in integrating the data after multivariate analysis qualitatively indicated that uMAIA was better at recapitulating the ground truth (**Figure S7a, b**).

We next asked to what extent each method enabled biologically meaningful comparative analysis between regions. We selected 5 regions annotated by the ABA that spanned multiple coronal sections (isocortex, cerebellum, hippocampal formation, hypothalamus, and thalamus) and applied a two-sided t-test between pairs of regions using average values across the sections. Comparing these results to the ground truth images allowed us to identify false-negative (FN) and false-positive (FP) detections (**Figure S7c**). uMAIA followed ground truth trends more often than other methods, resulting in a lower FP and FN count rate (FPR=0.07, FNR=0.12) than z-normalization (FPR=0.15, FNR=0.24) and ComBat (FPR=0.18, FNR=0.22), approaching the lower bound of FP and FN rates (FPR=0.05, FNR=0.08, see Methods 3.3) (**Figure S7d, e**).

To assess the ability of the methods to correct intensity distortions seen in real data, we applied them to the matched peaks from our zebrafish embryo dataset. In addition to ComBat, we tested a deep-learning method, scArches, that recombines features to integrate the data [44]. uMAIA normalization reduced the high-frequency intensity differences that were likely attributable to experimental noise in the foreground mode, while retaining signal mean levels (**Figures 4d, e**; and **S8a-d**). As there is no existing ground truth for this dataset, we rely on a multivariate and clustering analysis jointly applied to the pixels of all sections to evaluate performance, with the expectation to observe the same clusters in corresponding regions of consecutive sections if normalization was successful. We used ComBat, scArches, and uMAIA to normalize the 72 hpf zebrafish dataset and applied PCA followed by the K-means clustering algorithm [39, 44] (**Figure S8e,f**). Visualization of the clusters across all sections revealed that the grouped pixels after uMAIA normalization were most consistent across sections (**Figures 4f, g**; and **S8f**).

Application of uMAIA allows for the integration of MSI data for whole-organism and organ atlases, helping reveal patterns in lipids distributions and elucidate their roles.

### Emergence and diversification of metabolic territories during zebrafish development

Previous studies have shown that lipids undergo significant remodeling throughout zebrafish development [20, 21]. To gain a more comprehensive coverage of lipid species and clarify the extent of such changes between different developmental milestones, we performed bulk lipidomics by a maximum-coverage and high-throughput targeted HILIC–MS/MS workflow [45]. The selected time points comprised zebrafish embryos at four developmental stages: late gastrula (8 hpf), the latest time point of somitogenesis (24 hpf), pharyngula (48 hpf), and hatching (72 hpf) stages (**Figure 5a**) [46]. We quantitatively profiled 850 individual lipid species in zebrafish embryos, which revealed substantial lipid remodeling during development (**Figures 5b, c**; and **S9**). Specifically, hexosylceramides (HexCers) levels were found to increase the most, rising >10-fold from 8 to 72 hpf. Ceramides (Cers), sphingomyelins (SMs), phosphatidylserines (PSs), and phosphatidylglycines (PGs) also increased in their amounts, while phosphatidylcholines (PCs) and lyso-PCs (LPC)s significantly decreased (**Figure 5c**). Triacylglycerols (TGs), many of which reside in the yolk and serve as energy and building block storage for embryogenesis, did not collectively change in their levels (**Figures 5c**; and **S9a-c**). Surprisingly, the concentrations of TGs consisting of 54 or more carbon atoms in their fatty acid chains increased over development (**Figure S9d, e**).

**Figure 5:**
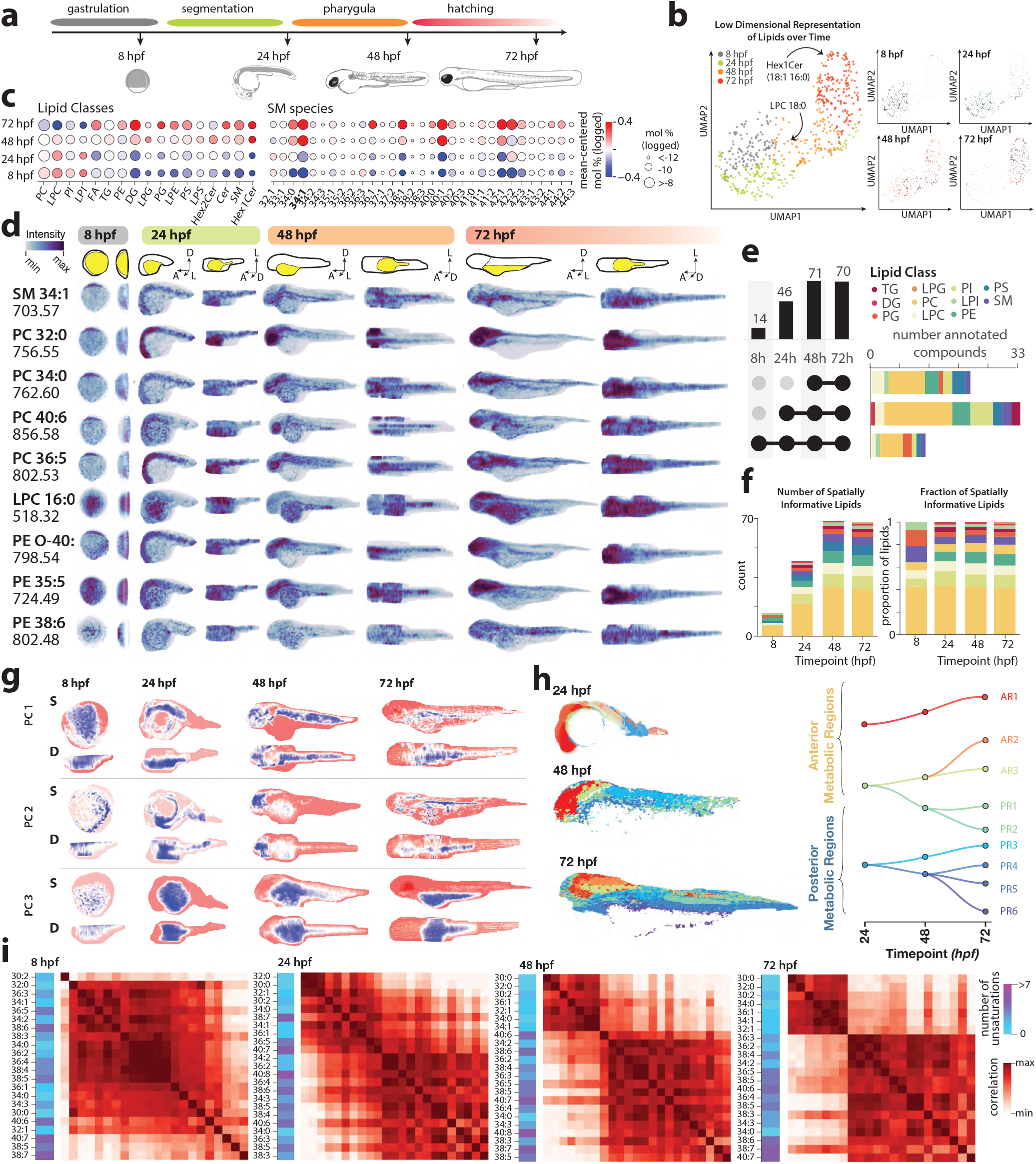
4D metabolic atlas of zebrafish development. **(a)** Overview of developmental stages used for bulk LC/MS and MALDI-MSI acquisitions with the main developmental stages indicated. **(b)** Low dimensional representation (UMAP) of bulk LC/MS data. Points represent lipids and are color-coded by the time point at which the concentration peaks (left), or relative changes in concentration over time (right). **(c)** Dot plot indicating overall lipid concentration changes and absolute quantities averaged over classes (left) and specific sphingomyelin species (right) for each developmental stage. **(d)** 4D distributions for selected lipids for selected time points (sagittal and dorsal projections, schematic of orientation shown in top row with yolk outlined in yellow; D: Dorsal; A: Anterior; L: Lateral.). Note: precise orientation of 8 hpf embryo is unknown. **(e)** Quantification of spatially-localized lipids across the sampled time points. Upset plot depicting total molecule counts for each timepoint (left). The timepoints that are considered within each set are indicated by solid black circles. Stacked bar plot depicting lipid class break up for different sets shown in upset plot (right). **(f)** Total number (left) and proportions (right) of spatially-informative molecules detected in specified time points stratified by lipid class. **(g)** Visualization of the first three principal components over each of the developmental time points, sagittal and dorsal views (upper rows). Component loading distribution per section displayed (bottom rows). **(h)** Pseudolineages and tissue clusters across sampled developmental time points split generally into anterior and posterior metabolic regions. **(i)** Correlation matrices between PC species for each developmental time plot. Colorbar indicates number of unsaturations for each lipid species.

However, bulk measurements cannot inform us of tissue-specific lipid compositional changes and thereby potentially overlook the importance of given lipids during tissue patterning. In order to properly fathom the spatial distributions of lipids, we built a 4D atlas. We considered our MALDI-MSI acquisitions comprising the selected timepoints and applied uMAIA to create a well-integrated dataset. Specifically, for each time point, we collected sagittal sections encompassing one whole embryo and additional representative sections as biological replicates. From this dataset we annotated by exact m/z matching 216 lipids that were also detected by bulk lipidomics and for which we could reconstruct 3D distributions (**Figures 5d**; and **S10a**). When inspecting these images, we observed that several lipids underwent a substantial spatial reorganization as development proceeded, localizing to specific structures by 48 or 72 hpf (**Figure 5d**).

We systematically assessed these changes and found that the number of lipids showing spatial patterns increased by threefold from 14 at 8 hpf to 46 by 24 hpf, leveling off for a total number of 70 species at 48 and 72 hpf stages (**Figures 5e, f**; and **S10b**, see Methods 4.1). The initial increase in localized lipids observed at 24 hpf aligns with the end of the segmentation phase when the primary organs and somites are established [46]. PCA analysis and clustering of pixels at each time point, using individual lipid levels as features, resulted in the definition of compositionally similar sets of pixels. These clusters of pixels were distributed within the 3D space of developing embryos in a spatially coherent manner, defining regions that we named *lipid territories*. Analysis of lipid territories across the different stages of development revealed an overall increase in the complexity of spatial lipid distributions (**Figure 5g, h**).

We connected metabolic territories between time points based on their similarity in lipid composition and observed that metabolically-related lipid territories occupied equivalent body regions at the different time points (**Figure 5h, left**). As complexity increases during development we set to follow the emergence of lipid territories by considering their pseudolineage, i.e., their connections over time, based on compositional similarity and investigated trends in pseudolineage branching (**Figure 5h**, right, see Methods 4.1). Notably, anterior metabolic regions did not appear to branch, suggesting that anterior tissues are already metabolically defined at the earliest stages of development considered as opposed to posterior tissues (**Figure 5h**). In support of this finding, anterior territories at earlier time points harbored more diverse metabolic content compared to those that resided posteriorly (**Figure S11a**, see Methods).

A lipid class that underwent substantial spatial reorganization during development is that of PCs. As time progressed, the emergence of a two-component structure in covariance between PC lipids became gradually evident. Intriguingly, PC species largely partitioned according to their degree of unsaturation (**Figure 5i**). PC species with lower numbers of double bonds were more localized to anterior tissues, constituting mainly the head, while the inverse was true for species with higher numbers of double bonds (**Figure S11c-e**).

Overall, this analysis indicates that lipid metabolic reactions are finely diversified and specialized in space and time during development. Biochemical remodeling and emergence of metabolic territories take place over time, with significantly more complexity present at 72 hpf compared to earlier time points. Importantly, this specialization revealed by spatial analysis was not deducible from bulk lipidomic experiments.

### Lipid-defined anatomy of a 72 hpf zebrafish embryo

The compelling finding that lipid metabolism is organized in space motivated us to directly investigate the correspondence between anatomical structures and biochemical organization. To do so, we focused on the most mature timepoint in our dataset, the 72 hpf zebrafish, as it harbored the greatest complexity in tissue composition as well as in spatial lipid distributions.

Annotation using LC-MS/MS-based lipidomic data allowed to identify and image 157 lipids (141 membrane lipids) across all inspected sections. This enabled the 3D imaging of approximately 20% of membrane lipid species, accounting for around 30% of the overall membrane lipid molar fraction (excluding cholesterol). The reconstructed lipids in the 72 hpf zebrafish embryo spanned various classes, including PC (51 species; 79.9 mol% of imaged lipids), LPC (15 species; 5 mol%), PE (15 species; 3.2 mol%), PS (22 species; 2.7 mol%), Sphingolipids (10 species; 4.7 mol%), TG (16 species; 0.04 mol%), PG (7 species; 0.06 mol%), PI (6 species; 1.7 mol%), LPI (3 species; 0.002 mol%), LPE (3 species; 0.01 mol%) and DG (9 species; 2.6 mol%) (**Figures S12, S13a-d**).

Manual inspection of these 3D lipid images revealed interesting spatial distributions. Remarkably, the localization of specific lipids faithfully delineated body parts such as yolk and yolk extension (TG 52:4), central nervous system (PC 34:2), hindbrain (PC 37:1), musculature and internal organs (PC 32:2) (**Figure 6a, b**).

**Figure 6:**
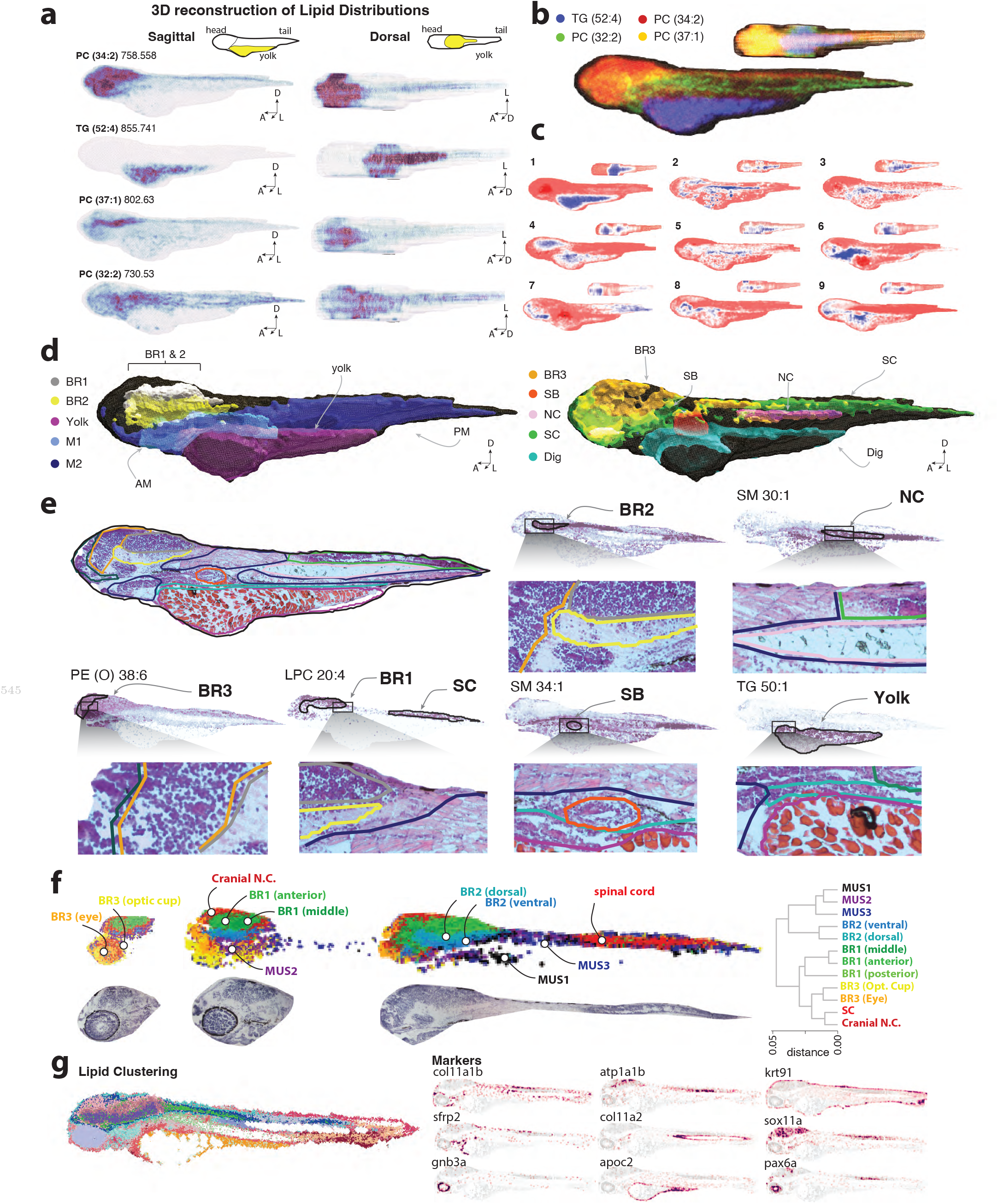
Characterization of metabolically-defined tissues in the 72 hpf embryonic zebrafish. **(a)** Sagittal and dorsal view of 3D reconstructions for 4 lipids with m/z value and lipid indicated. Schematic indicating yolk, head and tail displayed above. D: Dorsal; A: Anterior; L: Lateral. **(b)** Overlay of the 4 lipids visualized in (a). **(c)** Sagittal and dorsal views of the first 9 principal components retrieved after selecting for annotated lipids. **(d)** Visualization of metabolically-defined tissue clusters after application of unsupervised clustering algorithm (sagittal view); SC:spinal cord; NC:notochord; BR1, BR2, BR3: brain regions; SB=swim bladder; M1, M2=Musculature; Dig: digestive system. **(e)** Top row: medial section (sagittal view) of H&E staining overlayed with contours of metabolic regions shown in (d). Lipids that spatially localize to specific regions are indicated along with an inset of the H&E image. Both images are overlayed with contours representing delineated clusters. **(f)** Subclustering of neural regions (upper row) with corresponding H&E sections (bottom row). Dendrogram displaying relative distances between clusters (right). NC: neural crest, MUS: musculature **(g)** Lipid clustering of sagittal section of a 72 hpf (left) and marker gene distributions from a consecutive section from HybISS (right).

This correspondence between lipid distributions and anatomy generalized to other lipids, as revealed by the first 9 principal components, whose spatial representations unveiled more distinct anatomical distributions throughout the organism (**Figure 6c**). Moreover, a sorted distance matrix representing voxel similarity in lipid space displayed a block-like pattern suggesting the existence of a finite number of discrete lipid metabolic states in the embryo (**Figure S13e**). Clustering voxels into discrete categories (see Methods 4.2) resulted in lipid territories which, after comparison with hematoxylin and eosin (H&E) staining, identified several organs, such as the notochord, swim bladder, spinal cord and different areas of the brain (**Figure 6d, e**). To reveal if finer subdivisions could be discovered within lipid territories, we focused on the nervous system, as previous results suggested that it possessed high lipid diversity, and re-applied the clustering approach (**Figure S11a**). Interestingly, this subdivision tracked with anatomical structures: more subregions emerged after the second iteration of clustering that matched known tissues such as the eye and optic cup that differed mostly in their concentrations of LPC, PC and PG species (**Figures 6f** and **S13f,g**).

As a further test for the correspondence between lipid territories and anatomical structures, we performed Hybridization-based in situ sequencing (HybISS)[47] on a sagittal section of the 72 hpf zebrafish and used a consecutive section for MALDI imaging. A panel of tissue markers was used to highlight embryo anatomy (see Methods 4.3), while lipid-based pixel clustering was employed on MALDI-MSI data to retrieve lipid territories. As shown in **Figure 6g** several tissue markers faithfully localized to lipid territories, indicating that lipid distribution varies according to embryo anatomy and that embryo anatomy could be effectively derived from systematically assessing lipid distribution.

Prompted by the tight correspondence with tissue-marker gene expression we next investigated whether lipid territories could be simply derived from the expression of lipid metabolic genes. By analyzing publicly available single-cell expression profiles we found a limited match between genes implicated in metabolism and annotated cell identity due to the overall low expression of metabolism-related genes. Specifically, the 400 reported lipid metabolic genes (e.g., enzymes and transporters) were significantly less capable of predicting cell identity (R2 0.65; bootstrap estimate) than the 400 most variable genes (R2 0.78; bootstrap estimate), and not significantly different from the predictions given by a random set (R2 0.60; bootstrap estimate) [48]. (**Figure S14a, b**, see Methods 4.3). This result indicates that gene expression data may be inadequate to capture the relevant genes for predicting metabolic activity. We attempted to assess whether the localization of the metabolic genes, although weakly expressed, could still recapitulate anatomical features. We expanded our HybISS approach to include the most variable metabolic genes (see Methods 4.3). Apart from a few cases, metabolic genes were typically much noisier than marker genes and were less capable of identifying lipid territories compared to MALDI-MSI data (**Figure S14c-e**).

To biochemically characterize anatomical regions we performed an enrichment analysis of lipid species across the delineated lipid territories, identifying the one where each species was maximally abundant (**Figure 7a**). Stratifying these assignments by tissue revealed that several regions, including the swim bladder, digestive system, and regions of the brain, were predominantly characterized by a few lipid classes. This indicates that lipid species from the same class tend to localize within the same anatomical structures (**Figure S15a**).

**Figure 7:**
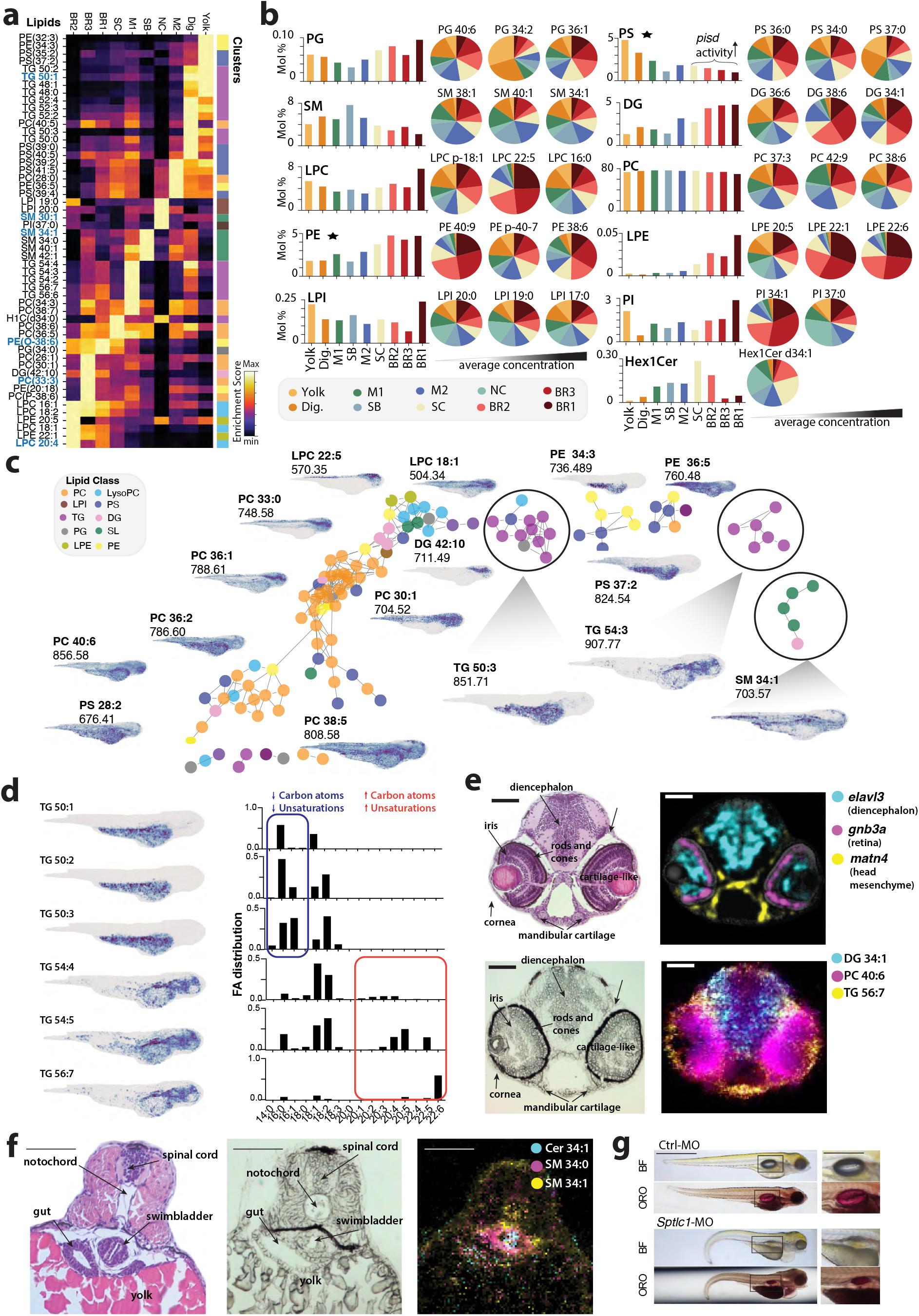
Biochemical organization of metabolically-defined tissues in the 72 hpf embryonic zebrafish. **(a)** Enrichment heatmap of lipids within clusters shown in Figure 6d. **(b)** Mol% composition of metabolite-defined clusters stratified by lipid class. Pie plots from left to right show 3 lipids of increasing concentrations (lowest, median, and maximum) and display localization according to mol%. **(c)** Force-directed layout representation of lipid-lipid co-localization. Lipid class indicated by colorcode. Images beside node clusters display representative lipid distribution of node clusters. **(d)** Subset of TG species images (left) with fatty acid compositions (right). Blue and red indicate fatty acids with low and high carbon content and unsaturations, respectively. **(e)** H&E (upper left), brightfield (lower left), genes from HybISS (upper right) and lipids from MALDI-MSI images overlaid (bottom right) from a coronal section through the head of a 72 hpf zebrafish. Scale bar 100um. **(f)** H&E (left), brightfield (middle) and lipids from MALDI-MSI images overlaid (right) from a coronal section through the swim bladder of a 72 hpf zebrafish. Scale bar 100um. **(g)** Brightfield (BF) and ORO stainings of control and *Sptlc1* mutant fish with zoom in of swim bladder in right column. Scale bar 1 mm.

By combining spatial and bulk lipidomics data, we obtained the relative concentrations between lipid classes. We found PC class lipids, as expected, constitute a significant proportion of lipids across all identified clusters, while other classes showed more variability in their concentrations. For example, a divergent trend between PE and PS concentrations was evident (**Figure 7b**) and may be ascribed to the expression of *Pisd*, which encodes a PS decarboxylase that converts PS to PE [49]. In accordance with *Pisd* being localized to the nervous system [48], PE concentrations were higher in neural tissues, while those of PS were reduced (**Figure S15b**).

Along similar lines, visualizing lipid-lipid relationships with a graph whose connectivity and layout is based on their spatial covariation revealed a strong biochemical organization by lipid class (**Figure 7c**). Thus, DGs and lysolipids populated different regions of the nervous system, TG species were observed to distribute between two biochemical clusters, SM species were colocalized to the swim bladder, spinal cord, and pericardial region, and PC species organized to two major lipid territories pertaining to the head or to more posterior tissues (**Figure 7c**).

Interestingly, TG species localization was generally dictated by fatty acid carbon chain lengths and unsaturation level. TGs with shorter chains (<54 C) and fewer double-bonds (<4) accumulated in the yolk, where they are known to play a role as energy reserves [20]. Surprisingly, TGs with longer chains and higher unsaturation levels populated anterior regions of the head and body yet distinct from the nervous system (**Figure 7c, d**). Previous studies in zebrafish embryos reported that neutral lipids are restricted to the yolk up to 24 hpf, while at later time points, they are also found in the head[20]. Our bulk lipidomics data and our spatial atlas consistently suggest that the TGs that populate extra-yolk anatomical districts are synthesized during development (**Figures S9d, e**) and contain longer >20C and more unsaturated fatty acids (**Figure 7d**). When the precise distribution of TGs in the head was investigated, we found that these localized to a mesenchymal compartment related to cartilage and bone *primordia* (**Figure 7e**) in line with the evidence that fish bones harbor extensive fat depots [50].

SM species localization to the swim bladder was also unexpected. The swim bladder is the organ responsible for regulating buoyancy in fish and is a homolog of the tetrapod lung [51–53]. It starts forming from the esophagus at 48 hpf and is inflated with air around 96 hpf. Similar to the tetrapod lung, the swim bladder inflation process requires the secretion of lipid-laden surfactant in the organ cavity to lower surface tension [52, 53]. SM started to accumulate in the prospective swim bladder at 48 hpf and became a distinguishable cluster by 72 hpf (**Figure 5d**) [54]. At this stage, the organ is organized into concentric tissue layers encompassing an inner epithelial layer, an intermediate layer of mesenchymal tissue, and an outer mesothelium [54]. When we inspected the distribution of SMs and related ceramides in coronal sections from 72 hpf embryos, we found that they populate concentric layers of the swim bladder, possibly reflecting the organ’s known tissue structure (**Figure 7f)**.

Intrigued by their peculiar distribution, we decided to perturb sphingolipid production to test its impact on embryo development. To this aim, we injected morpholinos targeting *Sptlc1* into embryos at the 1-cell stage (see Methods 1.5). *Sptlc1* is a key component of the enzymatic complex that catalyzes the first and rate-limiting step in sphingolipid biosynthesis [55], and evaluated the impact on how the embryo developed at 120 hpf. We observed that *Sptlc1* morpholino-injected embryos displayed developmental defects including tail curling and pericardial edema (**Figure S15c**). Among other defects, *Sptlc1* morpholino-injected embryos showed an impairment of their ability to inflate their swim bladder, as measured by a significantly different aspect ratio of the swim bladder (**Figure S15d**). Of note, a similar phenotype was episodically reported for an *Sptlc1*-KO line that, among other defects, also showed an inability to inflate their swim bladder [21]. Finally, Oil Red O staining [56] highlighted that *Sptlc1* morpholino-injected embryos were defective in their surfactant levels while they do not show histological alteration (**Figures 7g** and **S15e**). These data indicate that sphingolipids accumulate in the forming swim bladder during zebrafish embryogenesis and that they are involved in surfactant production and swim bladder inflation.

Altogether, this evidence reveals that during development, specific metabolic programs are established within different embryonic structures and are intimately related to their formation and function.

## Discussion

During embryogenesis, signals and spatially defined differentiation trajectories inform identical precursor cells to generate anatomical structures with distinct biochemical compositions. While morphogenic programs have been well-described through the lens of transcriptomics, understanding how metabolism evolves spatiotemporally in the context of embryogenesis is essential for comprehending the molecular processes that guide tissue pattern formation.

Here, we generated a 4D lipid metabolic atlas of vertebrate development to characterize the emergence of biochemical territories. To achieve this goal, we acquired a comprehensive dataset of high-resolution MS images from zebrafish embryos at different developmental stages. We also developed a general computational framework, uMAIA, which rethinks MSI data processing towards the integration of multiple acquisitions unlocking the potential for constructing whole-organism metabolomic atlases and enabling reliable cross-acquisition comparisons.

Previous MSI analysis tools have focused on the analysis of a single or a small number of acquisitions, yet the ability to coherently analyze data after the application of these tools drops quickly with dataset size, rendering analyses of >100 acquisitions virtually impossible. uMAIA marks a significant advancement in this sense, focusing on the creation of large information-rich metabolomic atlases, enhancing the depth and breadth of biochemical analysis, and introducing a level of scalability previously unattainable. Different acquisitions are made interoperable with conservative procedures and minimal explicit assumptions: uMAIA creates a normalized and coherent feature set, has no black-box components, and does not impute values or create dependencies between features. This advancement in data handling paves the way for studies that systematically combine spatial metabolomics with other modalities, such as spatial transcriptomics, proteomics, and ultrastructural data, unveiling more intricate dynamics within cells and tissues. We expect the integration of organismal-scale measurements of different biological dimensions will reveal heterogeneous regulatory circuits, as recently demonstrated by the finding of lipotypes [15].

In this work, we present a complete multivariate analysis of the MSI atlas we generated. Our results reveal a remarkable correspondence between lipid distributions and anatomical architectures, reflecting how metabolic programs are organized based on the cell type composition of different tissues. Importantly, transcriptomic methods showed suboptimal sensitivity to capture the spatial distributions of key lipid metabolic genes. This indicates that it is necessary to quantify lipids directly in order to address the spatial organisation of the lipidome.

When examining spatiotemporal lipid level changes over development, we encountered unexpected findings. We observed a peculiar patterning of PC species, whereby lipids with different unsaturation degrees are spatially segregated. As these lipids are quantitatively major membrane components and possess distinctive biophysical properties [57–59], their differential representation across various tissues is expected to impact membrane behavior. Determining how this pattern emerges mechanistically and understanding its physiological relevance are promising goals for future studies.

More generally, the colocalization of lipid classes to metabolic territories suggested that the territories are defined by the activity of metabolic pathways, and that they are possibly involved in specific tissue functions. Examples include the accumulation of sphingolipids in the swim bladder, where they might function as surfactant components, as well as long-chain TG species in bone *primordia* which may play a role in osteogenesis in the zebrafish head. We expect discoveries of this kind to help uncover links with disease and potential therapeutic targets, as exemplified by the aberrant completion of swim bladder organogenesis observed upon disruption of the sphingolipid biosynthetic pathway.

Global assessment of lipid distributions during embryo development revealed a close parallel between the unfolding of lipid metabolic territories and the process of organogenesis. Embryonic lipid territories did not specify simultaneously over time; the anterior regions, particularly the nervous system, exhibited the highest variation in metabolic composition and were the earliest to undergo fine metabolic patterning. Investigating whether this rostral specification of the lipidome is conserved or more pronounced in species with a finely regionalized central nervous system, such as mammals, is a promising area for future research. More generally, 3D and 4D atlases of multiple species will enhance our understanding of whether lipid patterning co-evolved with brain expansion and increase in complexity.

In conclusion, by developing uMAIA, we have made the unbiased analyses of large, organismal-level, MSI datasets tractable. Exploiting these new capabilities, we have conducted the first whole-embryo exploration of an important compositional aspect of living systems: their lipidome. Our results reveal significant heterogeneity along this underexplored axis, highlighting it as a compelling ground for discoveries in the regulation of development.

### Limitations

uMAIA remains bound to the limitations of MSI instruments, which capture only a fraction of all the molecular species in a sample, can alter the abundance estimation of species as a result of ion fragmentation, and depend on sample quality. Another limitation is related to how system requirements scale with dataset size: while uMAIA is thought to integrate and compare several acquisitions, the requirements of the method are expected to increase with dataset size. In settings where the co-analysis of hundreds of acquisitions and several millions of pixels is needed, the method will likely benefit from faster-compiled implementations or approximations to guarantee a manageable runtime.

In this work, we describe the changes in lipid distributions and the emergence and diversification of metabolic territories during zebrafish development. While metabolic precursors are stored in the yolk, we are not able to discern between the transport of inherited lipids from *in situ de-novo* synthesis during embryogenesis. Additional experiments including metabolic labeling approaches to reveal newly synthesized lipids in combination with spatial transcriptomic data and at higher temporal resolution would be necessary to address metabolic fluxes in space.

## Data and Code Availability

The raw data can be retrieved on METASPACE in .IBD and .imzML format at https://metaspace2020.eu/project/uMAIA. HybISS data are available at the following link: uMAIA is implemented in Python and available as an open-source package on GitHub at https://github.com/lamanno-epfl/uMAIA. Source code, installation instructions and tutorials are also available on GitHub.

## Author Contributions

H.H.S. developed the idea, designed and implemented the model framework, ran simulation experiments and performed corresponding analysis, analyzed the MSI and LC/MS data, created the figures and wrote the manuscript. L.A. developed the idea, collected and acquired all MSI zebrafish data, collected zebrafish embryos for LC/MS experiments, analyzed scRNASeq data, helped analyze MSI and LC/MS data, performed and analyzed HybISS experiments and wrote the manuscript. L.A.R., G.V. and C.J. collected zebrafish embryos and helped with biological interpretation of the data. G.V. performed morpholino injections. A.G. refined the normalization component of the framework for faster implementation. A.H. performed HybISS experiments and analysis. A.S. adapted HybISS pipeline for zebrafish, designed the probes and performed HybISS experiments. A.D.M. performed image analysis on raw HybISS acquisitions. J.P.M. performed initial LC/MS experiments. L.C. provided initial MSI data used for developing the framework, helped in their interpretation and advised on technical aspects. I.K. contributed to initial development of the pipeline for MSI data analysis. G.D.A and G.L.M. developed the idea, supervised the project, and wrote the manuscript. All coauthors read and approved the manuscript.

## Acknowledgements

This project has been made possible thanks to SNF grant numbers TMSGI3_218393, IZSEZ0_213427, 316030_183503, 310030_184926, 310030_219382, as well as support from the Kristian Gerhard Jebsen Foundation, the Swiss Data Science Center (SDSC) and SERI-funded ERC Starting Grant for Lipo-Trace. We thank members of the La Manno, D’Angelo, Oates and Gisou van der Goot labs for their generous feedback and discussions on the project, particularly Agostina Crotta Asis, Jean Maillat, Daria Korotkova. We also thank the entire EPFL Histology Facility for their assistance with experiments.

## SUPPLEMENTARY FIGURES

**Figure S1:**
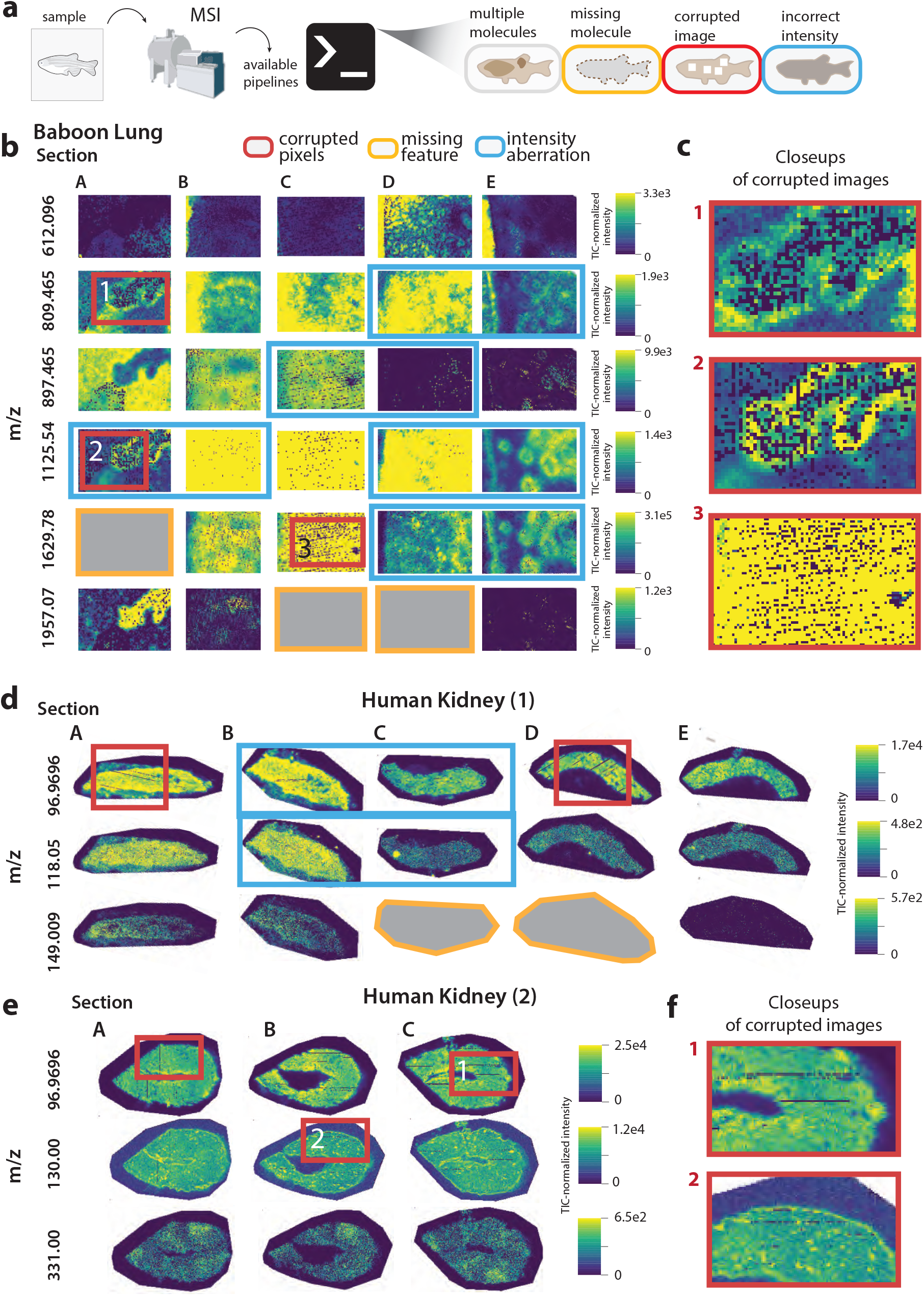
Showcasing different artifacts and data limitations impairing the analysis of MSI datasets. **(a)** Schematic illustrating the fact that MALDI-MSI data pipelines can produce outputs with artifacts and inconsistencies that can impair the downstream analyses of a combined dataset. **(b)** Represenatative MSI images of metabolites (m/z in bold) from public MSI datasets of the Baboon lung. Images were extracted from METASPACE. Different kinds of artifacts and limitations of the data that impact downstream analyses are highlighted in different colors. **(c)** Close-ups of MALDI acquisitions in (b) showing a scattered-like pattern of dropouts corrupting MSI images. **(d)** Representative MSI images from the Human Kidney. **(e)** Another set of MSI images from the Human Kidney. **(f)** Close-ups of the corrupted MALDI images in (e).

**Figure S2:**
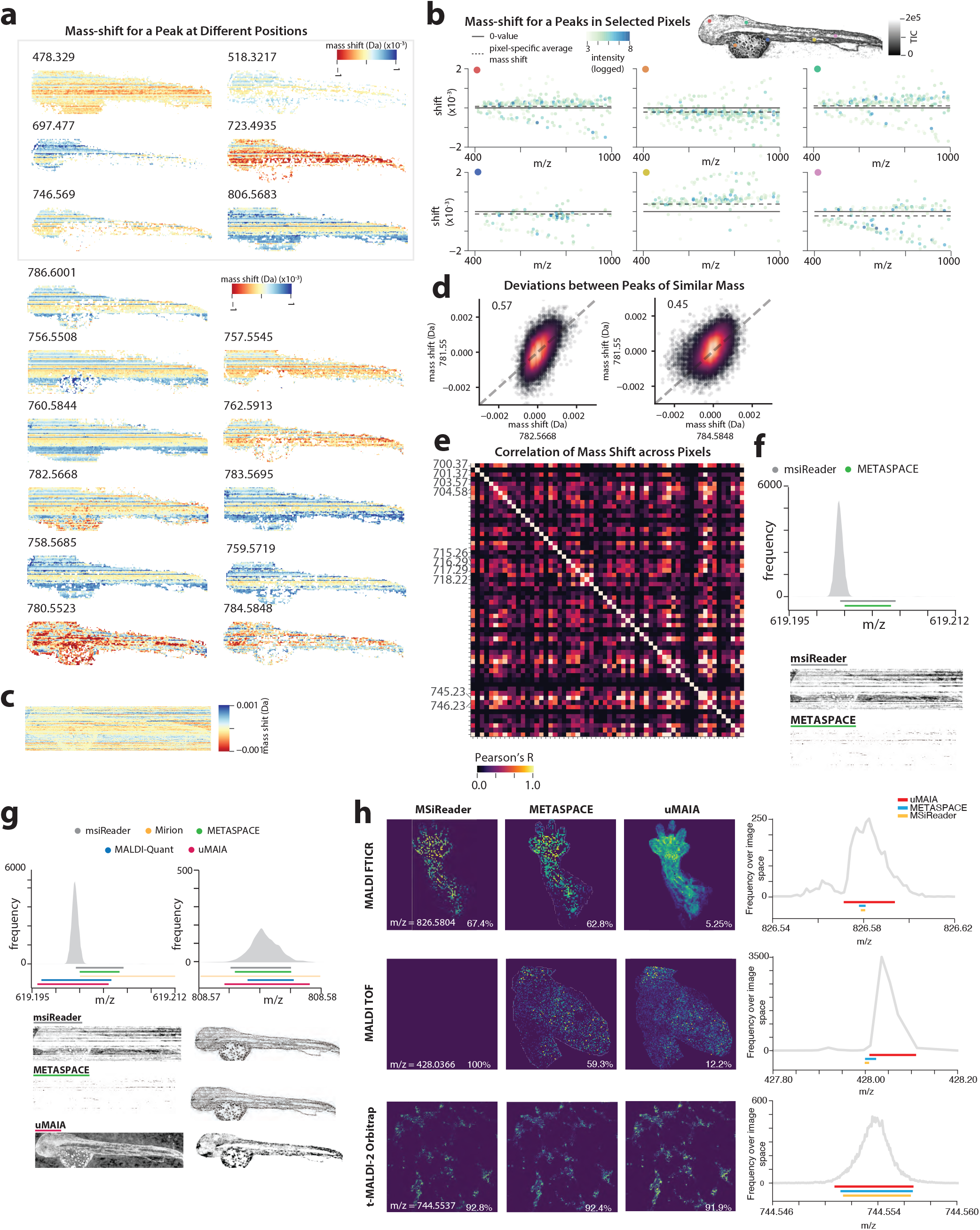
Characteristics of mass shifts and imaging of peaks by various software. **(a)**, Mass shifts represented as images for a variety of peaks across spectra (pixels). **(b)**, Mass shifts for all peaks within selected spectra indicated by color of dots (refer to image on top right for positioning) shaded by logged intensity. Solid and dotted lines represent 0 and average shifts within the spectrum, respectively. **(c)**, Overall average mass shift for all peaks. **(d)**, Scatterplots of mass shifts between quasi-isobaric molecules. Pearson’s R indicated in the top left hand corner. **(e)**, Correlation matrix of mass shifts between molecules with m/z values between 700-750. **(f)**, Histogram representation of spectra for specific peaks using non-adaptive binning methods (msiReader and METASPACE) with the called bin shown as solid lines (above) and resultant image (below). **(g)**, Histogram representations of spectra for two peaks, with identified bin from various methods shown as solid colored line (above) and correspondent image (below). **(h)**, Images for various MSI datasets using msiReader, METASPACE and uMAIA (left). Imaged peak m/z indicated on bottom left. The percentage of missing pixels indicated at the bottom of the image. The correspondent histogram representation is shown on the right with identified bin from various methods shown as solid colored line (right).

**Figure S3:**
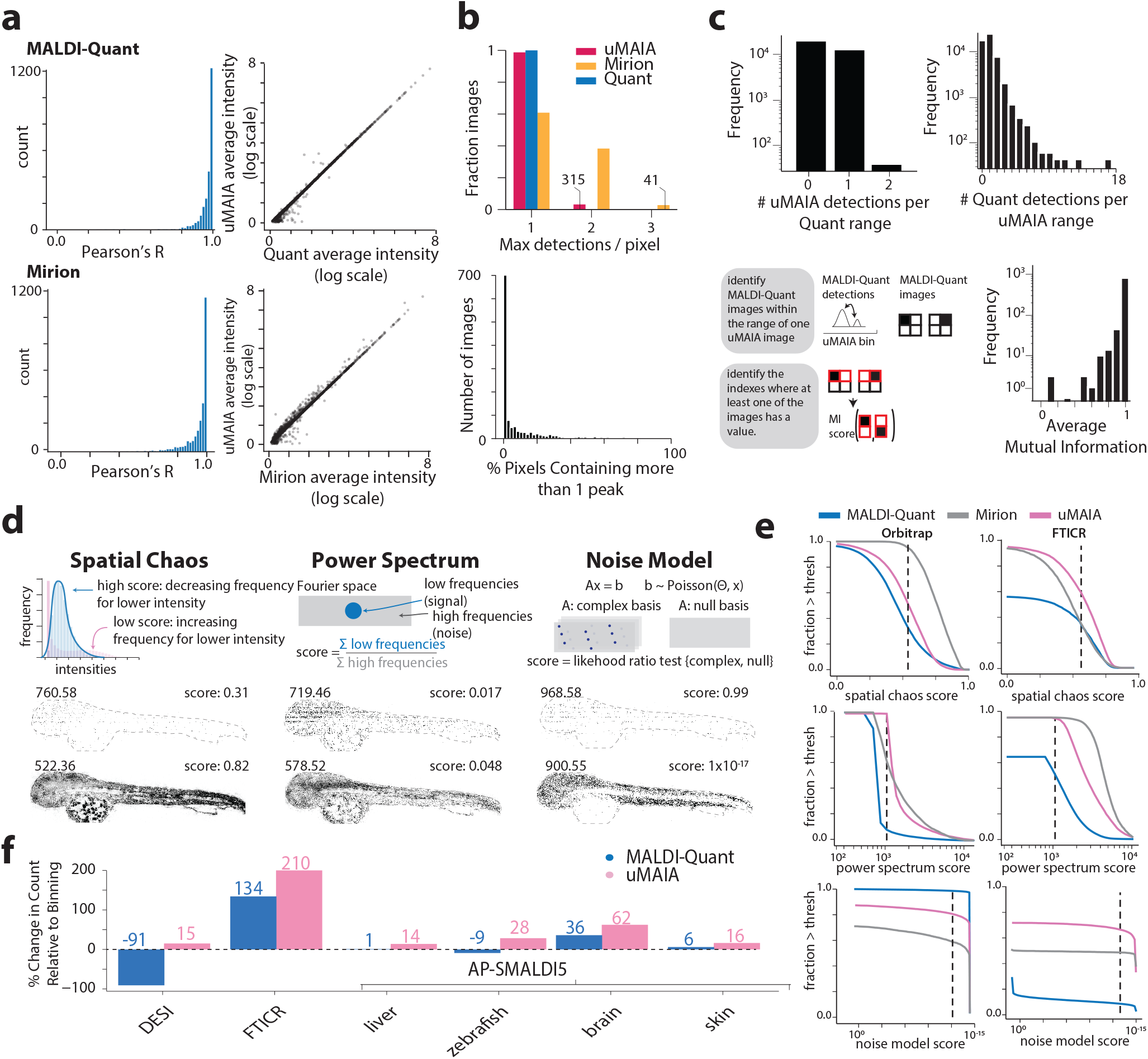
uMAIA peak-calling performance on MSI datasets. **(a)**, On the left, histograms of the similarities between images extracted for different compounds (MALDI vs uMAIA comparison above, and Mirion vs uMAIA below) Pearson’s R is computed on the intensity values. On the right, scatter plots of the average intensity in images retrieved. **(b)**, Fraction of images containing aggregated peaks for uMAIA, Mirion and MALDI-Quant (left). Histogram across images reporting on the proportion of pixels containing more than one peak from uMAIA-retrieved images for a single acquisition (right). **(c)**, uMAIA-retrieved bins were paired with overlapping MALDI-Quant-retrieved bins. For each MALDI-Quant bin, the number of uMAIA bins was quantified (upper left). For each uMAIA bin, the number of MALDI-Quant bins was quantified (upper right). In instances where multiple MALDI-Quant bins were detected in a given uMAIA bin, the average mutual information between images was calculated and plotted in a histogram (bottom row). **(d)**, Metrics used to quantify image quality: Spatial chaos, Power spectrum, and Noise model. Schematic depiction of metric (above) and example image with score displayed (below). **(e)**, Fraction of images (y-axis) surpassing Spatial chaos, Power spectrum, and Noise model metrics (x-axis) for uMAIA, MALDI-Quant and Mirion methods shown as a function of the metric. Left column: Orbitrap MSI data; Right column: FTICR MSI data. **(f)**, Percent change in count of images retrieved by uMAIA and MALDI-Quant surpassing Spatial chaos, Power spectrum, and Noise metrics with respect to the number retrieved by binning for various MSI acquisitions.

**Figure S4:**
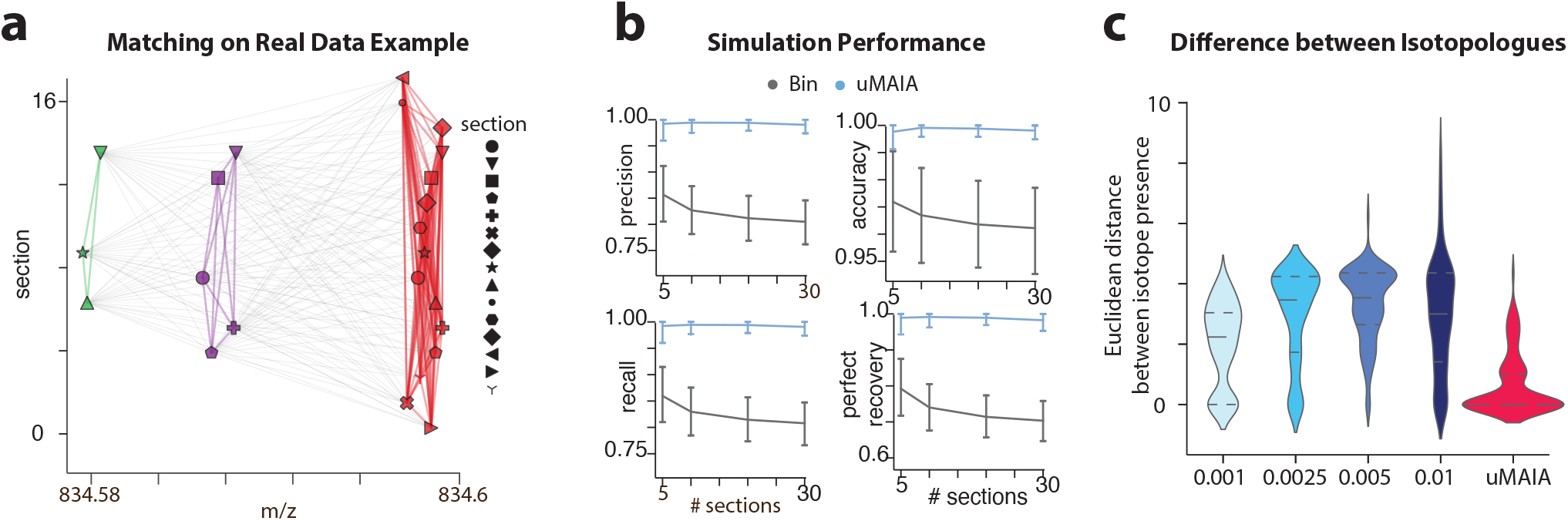
Characterizations of uMAIA’s molecule matching outputs. **(a)**, Diagram visualizing molecule matching outputs for a representative set of molecules. Allowed matches that are considered by the algorithm are represented with gray edges. Grouped molecules are colored by distinct colors and connected by colored lines. **(b)**, Results of a sensitivity analysis of molecule matching using different numbers of MSI acquisitions. Lineplots track different performance metrics (precision, accuracy, recall and perfect recovery) for simulated data. **(c)**, Violin plots reporting the distribution of Euclidean distances between isotopologue M+0 and M+1 presence across the acquisitions after featurization. Distributions are computed across all the pairs of sections for the 20 molecules with the highest signal intensity.

**Figure S5:**
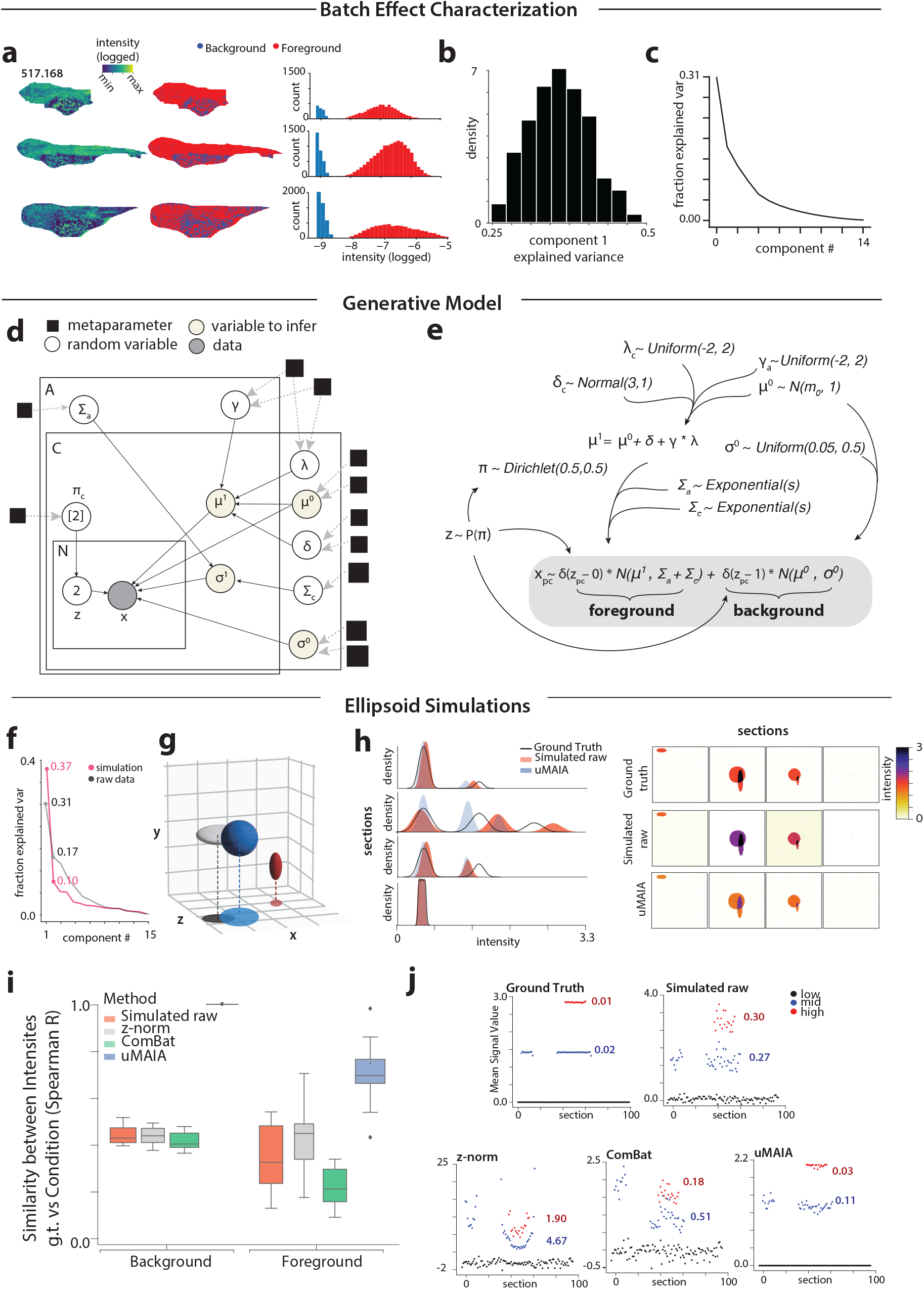
Batch effect characterization and evaluation of ground truth retrieval by uMAIA normalization. **(a)**, MSI images for m/z 517.17 at different mediolateral position (left) and corresponding background/foreground assignments (center) and histograms of intensity values (right), histogram bars are colored by the assignment (e.g. foreground or background). **(b)**, Histogram displaying the distribution of bootstrap estimates of the first eigenvalue of the empirical batch-effect matrix (See Methods). **(c)**, Line plot showing fraction of explained variance contributed by each principal component of the batch effects estimator (see Methods). **(d)**, Hierarchical Bayesian model in plate notation including metaparameters, latent variables and data for the estimation of the intensity-distorted signal. **(e)**, Generative model representation of the model in (c). **(f)**, Fraction of explained variance contributed by each component from SVD as in (c) (black line) overlaid with those from simulation data (red line). **(g)**, Visualization of spheres used in simulation. **(h)**, Intensity distributions in simulated data, comparing the ground truth, simulated raw and uMAIA normalized data for 4 sections (left) with corresponding images shown (right). **(i)**, Box and whiskers plots quantifying the similarity of the corrected images with ground truth results are computed across 4 sections and all variables of the simulation. Similarity is measured by Spearman’s R between intensities. **(j)**, Value of modes (low-, mid- and high-intensity) across sections for ground truth, simulated raw, z-normalized, ComBat and uMAIA corrected data for a single molecule (red: high intensity; blue:mid intensity; black: low intensity). Variance for the three categories is indicated by colored numbers.

**Figure S6:**
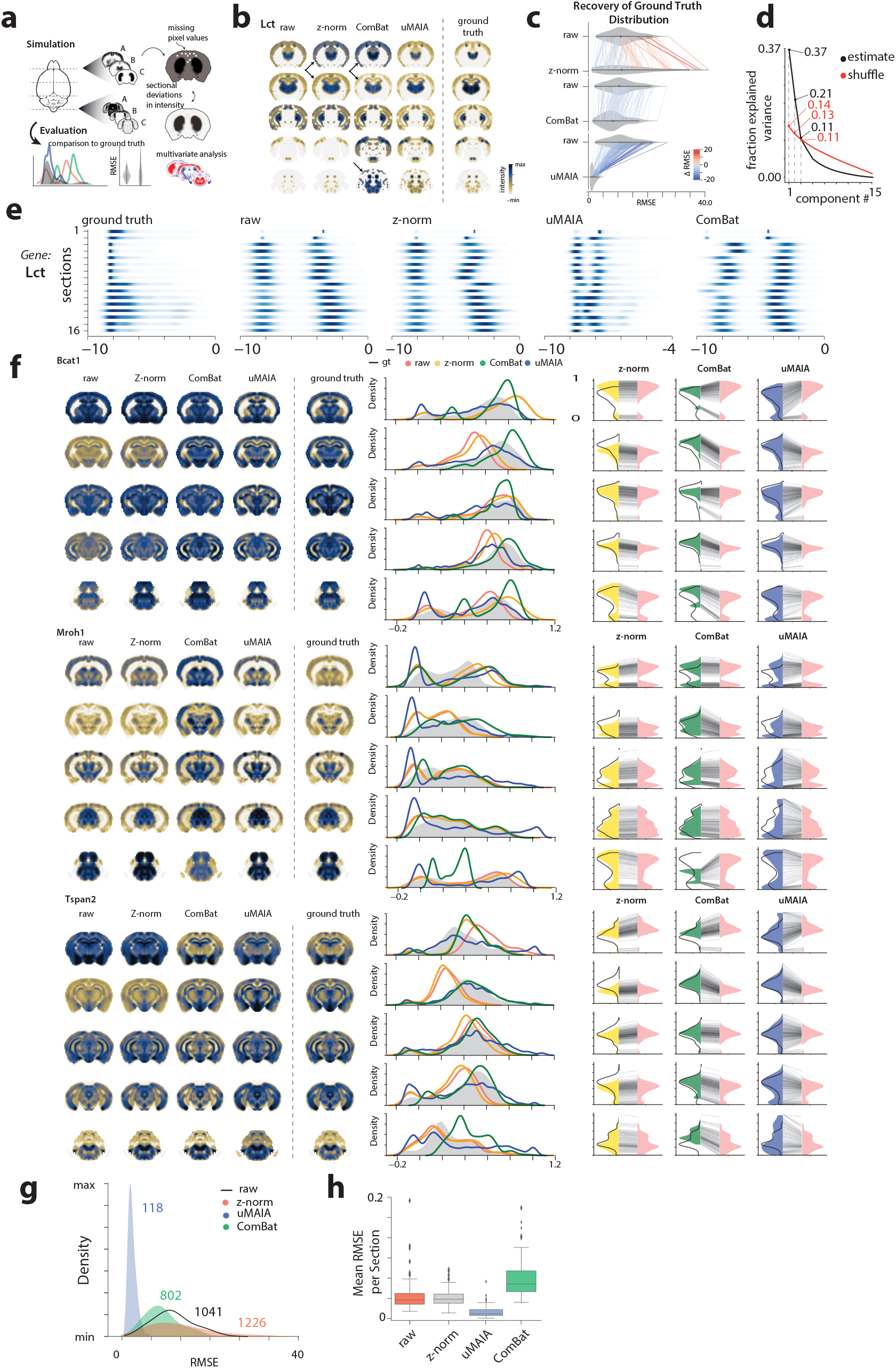
Comparison of simulated data batch effect correction using different methods. **(a)**, Schematic of simulation with ISH data from ABA and evaluation. **(b)**, Visualization of images corresponding to gene *Lct* of ground truth (right) and simulated raw data and data corrected with z-normalization, ComBat and uMAIA (left). **(c)**, Root Mean Square Error (RMSE) distribution with respect to ground truth images for simulated raw data and data corrected with z-normalization, ComBat and uMAIA. Lines are drawn between genes and colored by relative increase or decrease of RMSE compared to simulated raw data. **(d)**, Fraction of explained variance contributed by each component from SVD applied to batch effect estimates for ISH simulated data (black line) overlaid with those from shuffled data (red line). **(e)**, Intensity distribution densities across sections for gene *Lct* for ground truth, simulated raw, z-normalized, ComBat-normalized and uMAIA-normalized data. **(f)**, Images of genes *Bcat1, Mroh1, Tspan2* in ground truth, simulated raw, Z-normalized, Combat-normalized and uMAIA-normalized data (left) with corresponding intensity distributions (middle). RMSE plots for data corrected by z-normalization, ComBat and uMAIA, with comparison made against simulated raw data (pink distribution) and genes linked together by gray lines (right). **(g)**, Density of RMSE distributions for simulated raw data (black line) and data corrected with z-normalization, uMAIA and ComBat. Number displayed above distributions in affiliated color indicate total RMSE. **(h)**, Mean RMSE per section, for simulated raw data and data corrected with z-normalization, uMAIA and ComBat.

**Figure S7:**
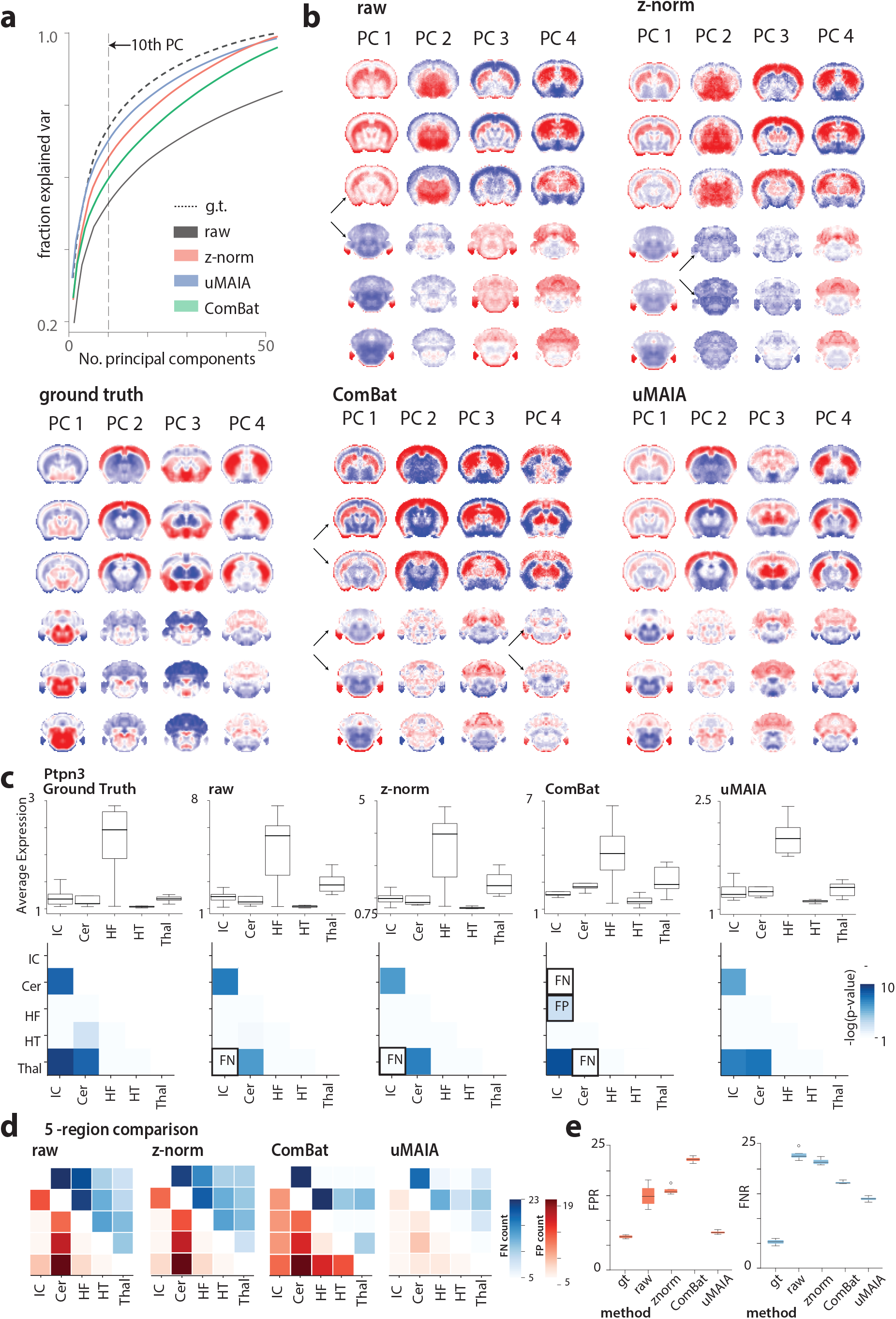
Evaluation of the rescue and preservation of ground truth signal after normalization using ABA-based MSI simulated data. **(a)**, Line plot tracking the fraction of explained variance explained by different numbers of principal components for ground truth, simulated raw data, and normalization on the ABA-based simulation (see Methods). Dotted line indicates the 10th principal component. **(b)**, Visualization of principal components for ground truth, simulated raw data and data corrected with z-normalization, ComBat and uMAIA. Black arrows indicate sections where significant batch effects still exist. **(c)**, Average expression of gene *Ptpn3* in 5 major anatomical regions. Box plots indicate variability of expression across sections for each setting: ground truth, simulated raw data and data corrected with z-normalization, ComBat and uMAIA (above). Heatmaps display the -log(p-value) of t-tests between any pair of regions that can further be used to distinguish false negative and false positive detections with respect to ground truth (below). **(d)**, Total number of false positives and false negatives after differential expression tests across all genes for 5 major anatomical regions. Heatmap is shown for simulated raw data and data corrected with z-normalization, ComBat and uMAIA. **(e)**, False positive and false negatives rates after differential expression tests using bootstrapped sections. Distributions shown for ground truth (gt), simulated raw and data corrected with z-normalization, ComBat and uMAIA.

**Figure S8:**
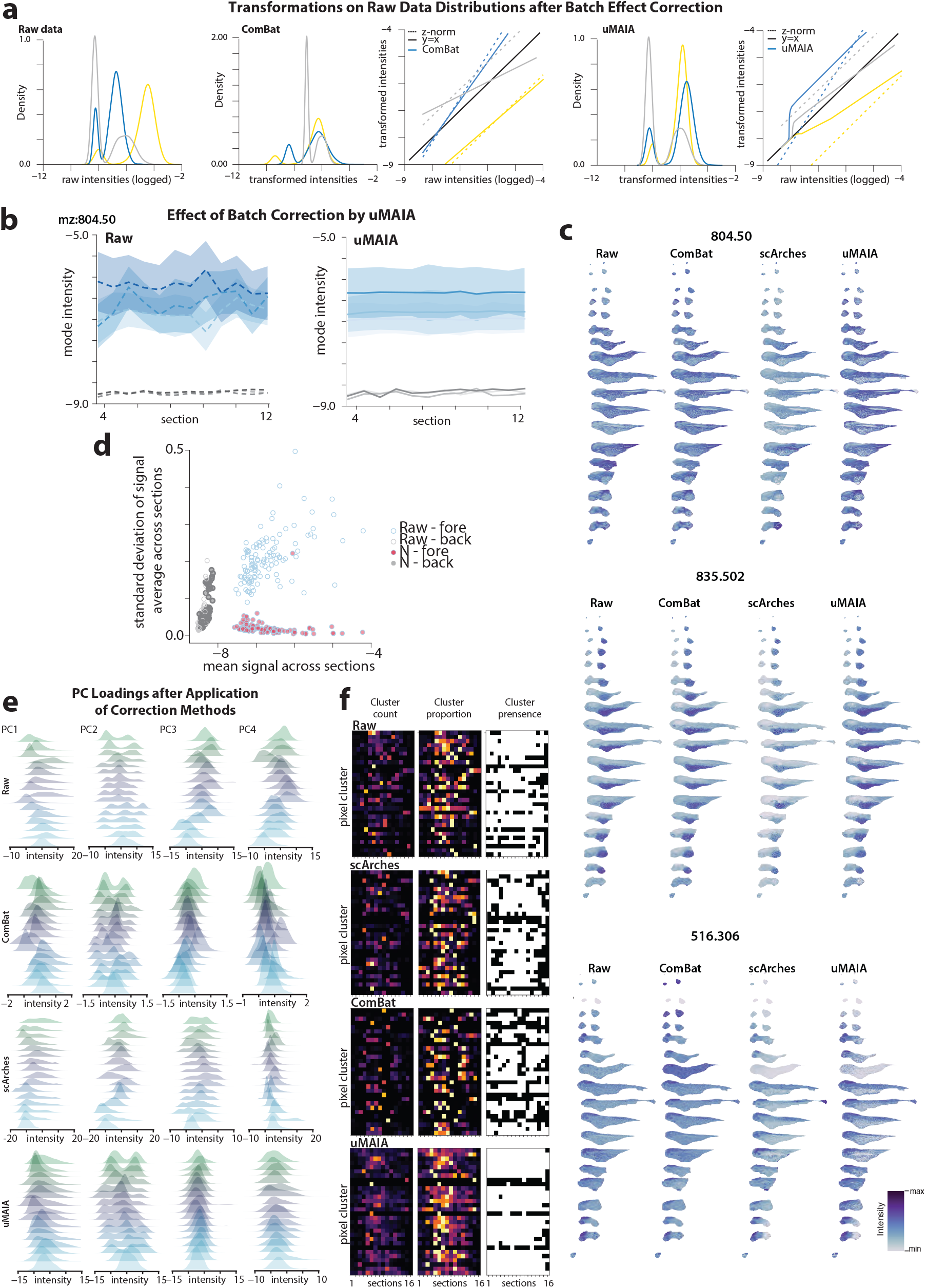
Comparison of batch effect correction methods on data. **(a)**, Signal intensity distributions across three acquisitions, color corresponding to the acquisition. On the left is the raw MALDI MSI data. To the right, the same data corrected with ComBat and uMAIA with related quantile-quantile plots representing the transformation between raw and normalized intensities. **(b)**, Line plot tracking the mean molecule (mz=804.50) intensity and variance (shaded area) in background and foreground modes across consecutive samples for raw and uMAIA-corrected data. **(c)**, Images of 3 peaks (mz=804.50, 516.30, 835.50) with spatially variable distributions for raw data and data corrected with ComBat, scArches and uMAIA. **(d)**, Scatter plot of mean signal vs standard deviation of signal average across sections for background and foreground modes for raw and uMAIA-corrected data (right) **(e)**, Distribution plots showing principal component loadings of pixels across different acquisitions for raw data and normalized data. The principal components appear to shift as a result of the distortion of many individual features, the normalization rescues their consistency between acquisition. **(f)**, Heatmaps representing the number, proportion and presence of pixel clusters across sections (‘cluster count’, ‘cluster proportion’ and ‘cluster presence’) for raw and corrected data.

**Figure S9:**
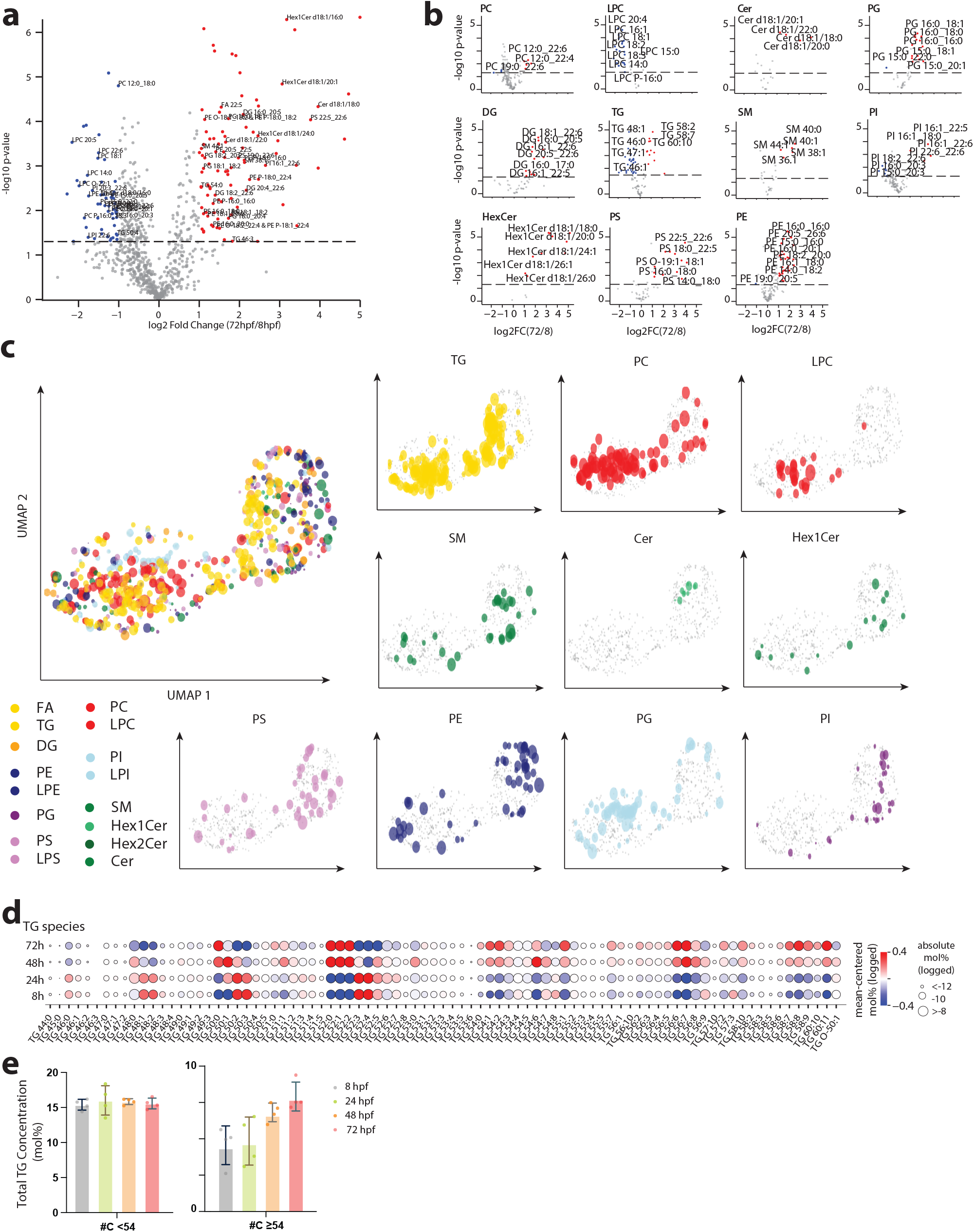
Lipidomic profiles over development measured by quantitative LC/MS. **(a)**, Volcano plots displaying fold change between lipids detected in 72 hpf and 8 hpf zebrafish from quantitative LC/MS data. **(b)**, Volcano plots as in (a) stratified by lipid class. **(c)**, Low dimensional representation (UMAP) of lipids abundances measured by quantitative LC/MS over developmental time color-coded by lipid class. Dot-size proportional to the log of measured concentration from LC/MS for each lipid species. **(d)**, Relative changes of TG species from bulk lipidomics across the time points. **(e)**, Total TG concentrations over time points stratified by total carbon content.

**Figure S10:**
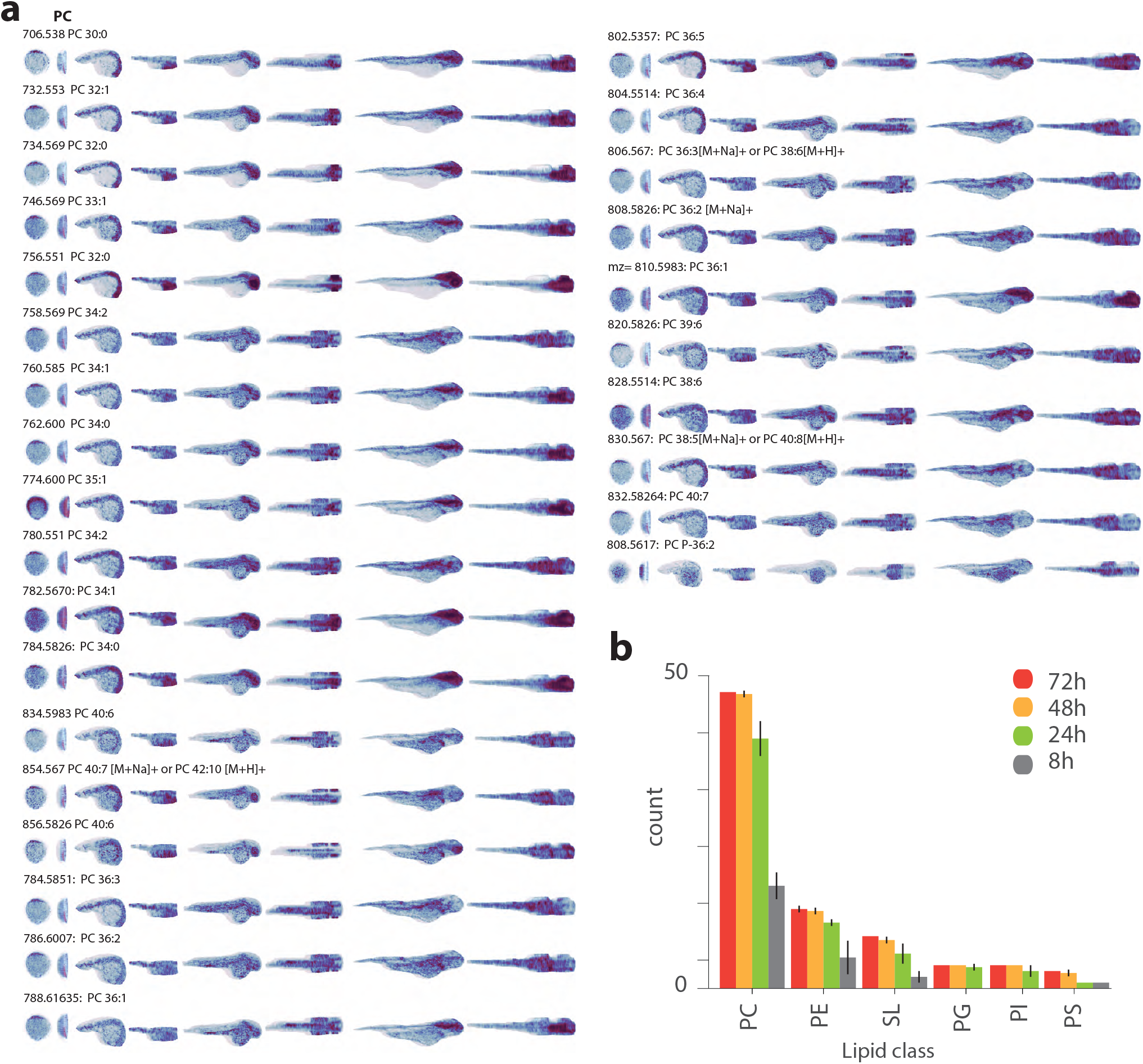
3D visualization of lipids during zebrafish development. **(a)**, Sagittal and dorsal views of 3D reconstructed images of lipids detected in MALDI-MSI data at 8, 24, 48 and 72 hpf sorted by lipid class. Note orientation of 8 hpf embryo is of uncertain orientation. Species name and m/z displayed on top left. PC: phosphatidylcholine; LPC:lysophosphatidylcholine; SL: sphingolipid; TG: triacylglycerol; PS: phosphatidylserine; PE: phosphatidylethanolamine; PI: phosphatidylinositol; LPI: lysophosphatidylinositol; PG: phosphatidylglycerol; LPE: lysophosphatidylethanolamine; DG: diacylglycerol. **(b)**, Number of spatially informative lipids stratified by class at 8, 24, 48 and 72 hpf time points of embryonic zebrafish development. Error bars represent standard deviation between counts across replicates (n=2).

**Figure S11:**
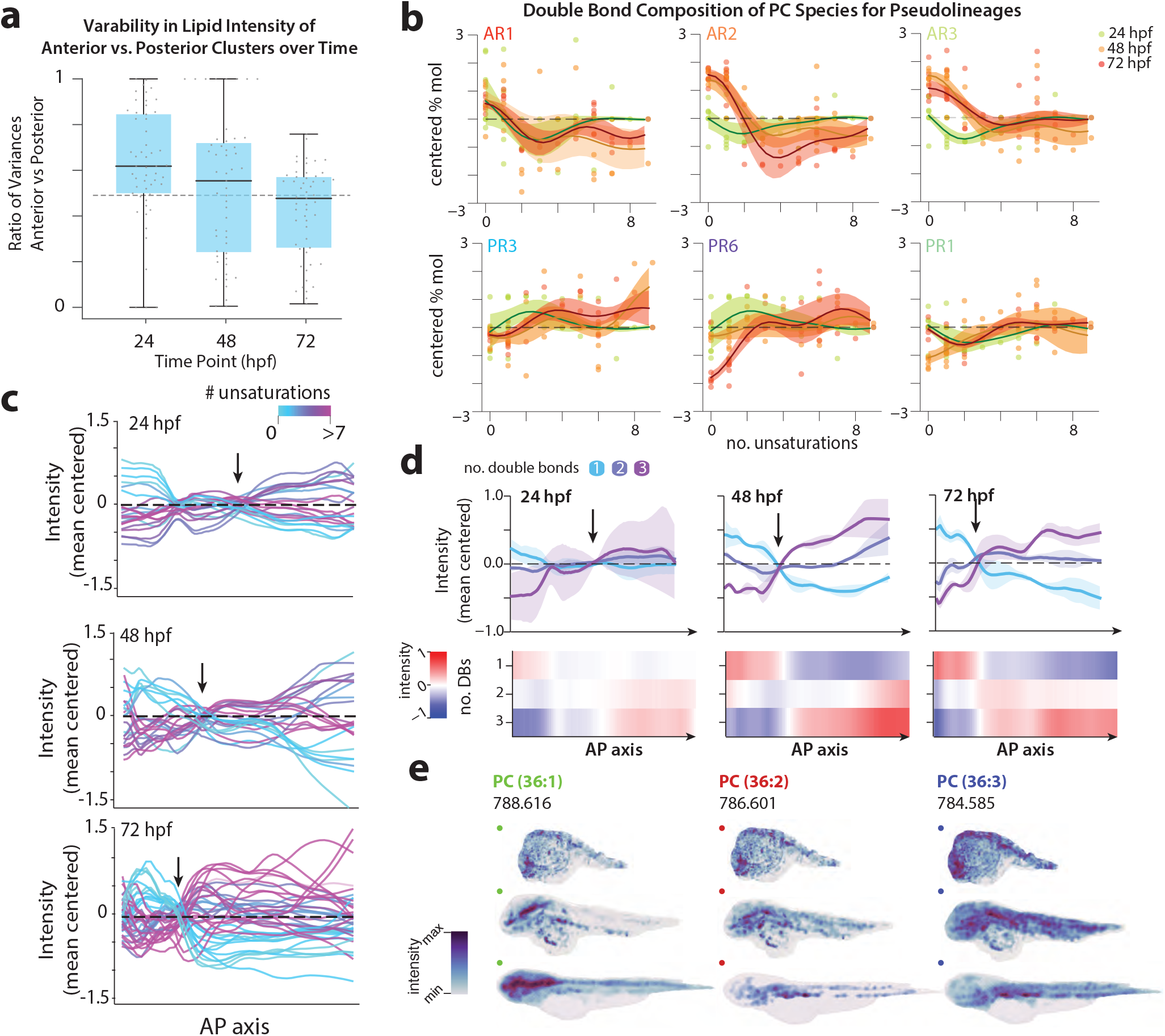
4D MSI atlas of zebrafish development. **(a)**, Box and whiskers plot displaying the distribution variability ratio between different metabolic regions, plot is stratified by time point, each dot represents a lipid. **(b)**, Mean-centered mol% of PC lipids for different pseudolineages as a function of their unsaturation content. Green: 24 hpf; Orange: 48 hpf; Red: 72 hpf. **(c)**, Lipid intensity trends for PC species along the AP axis for each developmental stage. Varying degrees of unsaturations are indicated by colors. Heatmap of mean-centered average lipid intensity for the 3 species over developmental time (lower row). **(d)**, Visualization of sagittal view of PC 36:1, PC 36:2 and PC 36:3 shown in (e).

**Figure S12:**
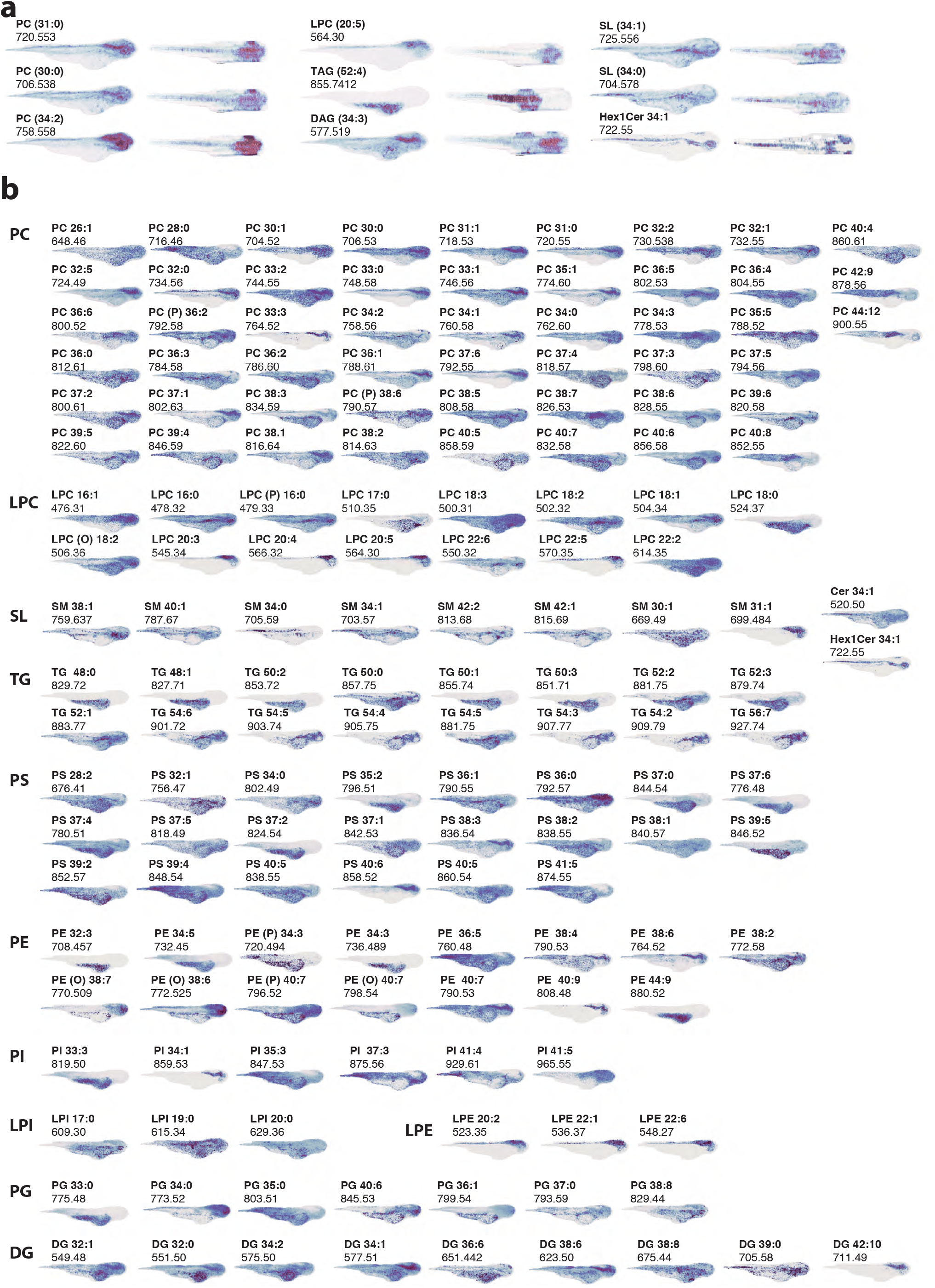
3D visualization of lipid distributions across the 72 hpf zebrafish embryo. **(a)**, Lateral and dorsal views of 3D distributions of lipids at 72 hpf. Intensities across the whole volume are presented using maximum intensity projection visualization. **(b)**, Lateral views of 3D distributions of lipids detected in MALDI-MSI data at 72 hpf sorted by lipid class. Species name and m/z displayed on top right. PC: phosphatidylcholine; LPC:lysophosphatidylcholine; SL: sphingolipid; TG: triacylglycerol; PS: phosphatidylserine; PE: phosphatidylethanolamine; PI: phosphatidylinositol; LPI: lysophosphatidylinositol; PG: phosphatidylglycerol; LPE: lysophosphatidylethanolamine; DG: diacylglyceride

**Figure S13:**
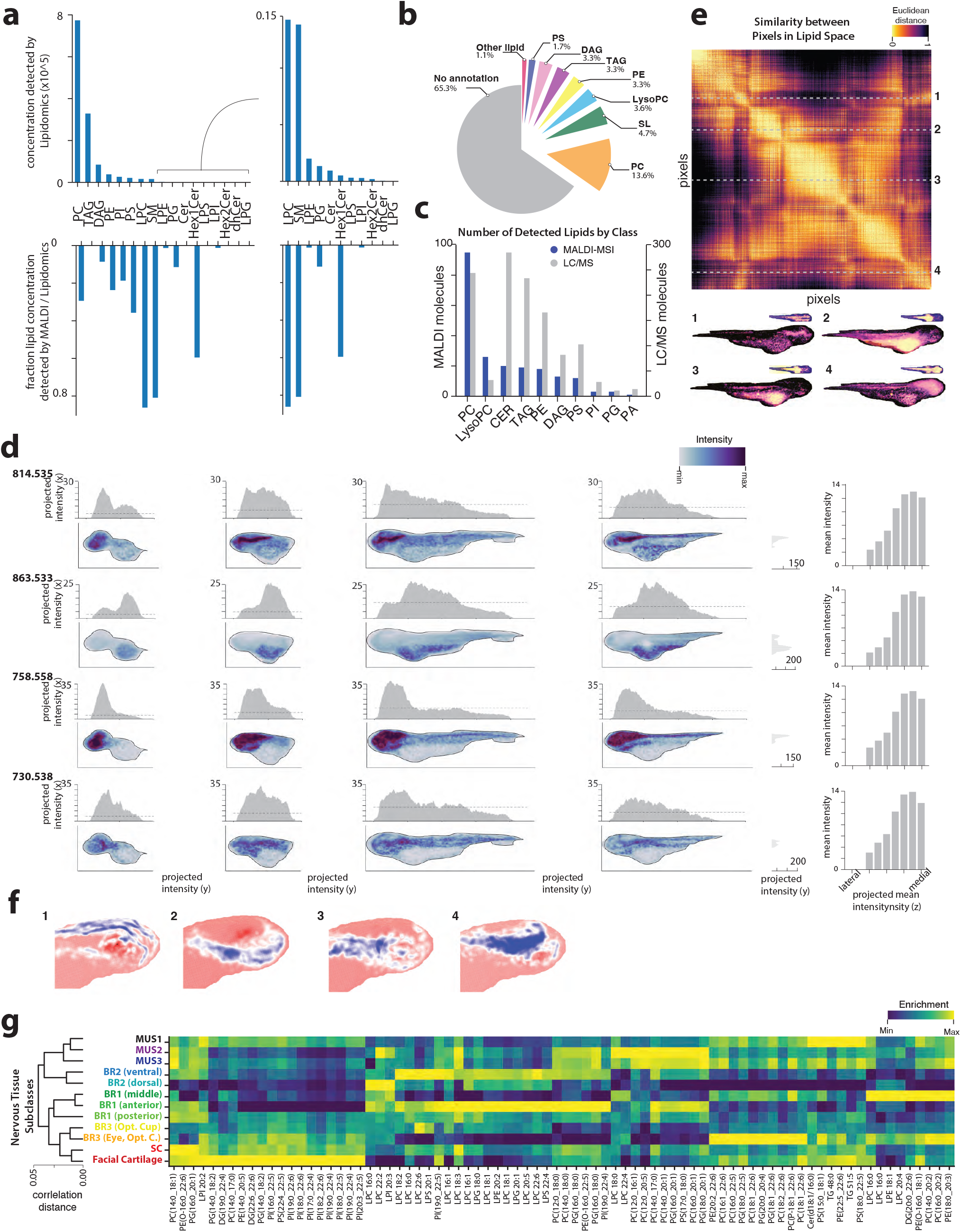
Lipid distributions and variability across 3 dimensions for the 72 hpf zebrafish embryo. **(a)**, Barplots show the concentration of different lipid classes as estimated by a quantitative LC/MS experiment (top row). Relative intensity values of MALDI-MSI are shown below for comparison (bottom row). **(b)**, Proportion of lipid species detected by MALDI-MSI stratified by class. **(c)**, Barplot reporting the number of lipids detected by MALDI-MSI compared to LC/MS stratified by lipid class. **(d)**, Visualization of the distribution of 4 lipids (rows, mz= 814.535, 863.533, 758.558, 730.538) in a set of four sagittal sections spanning the mediolateral axis. Histograms on the margins show the projected intensities across the x, y, and z axes. **(e)**, Heatmap of pairwise similarities between pixels lipid composition. The similarity metric used is Euclidean distance. On the left, the values of 4 rows of the matrix were visualized in correspondence with each pixel location to highlight the correspondence of the main blocks of the matrix with anatomical structures. **(f)**, Closeup of the head colored by the loading of the top four principal components. Images are 3d visualization using maximum intensity projection. **(g)**, Heatmap depicting lipid enrichment in different sub-brain region clusters. All plots in the figure refer to the 72 hpf zebrafish embryo. Rows are sorted as indicated in the dendrogram on the right.

**Figure S14:**
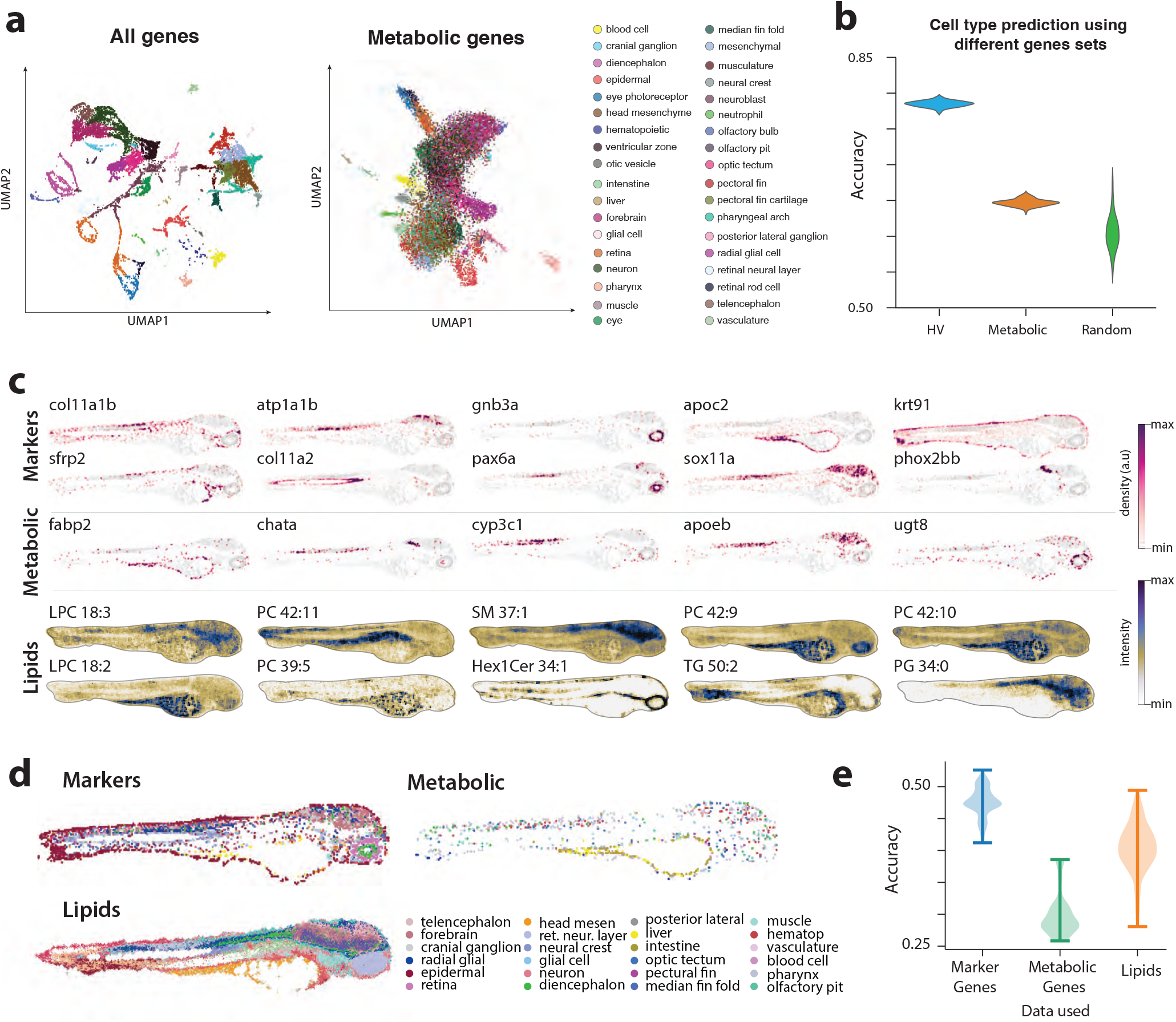
Comparison between spatial lipidomics and transcriptome of the 72 hpf zebrafish embryo. **(a)**, Low dimensional representation embeddings (UMAP) of cells’ gene expression from scRNAseq data [48]. Two embeddings are shown using either all measured genes or the subset that are associated with lipid metabolic processes. Cells are color-coded according to annotated cluster identity. **(b)**, Accuracy of models using different gene sets (Highly Variable: HV, Metabolic, and a random set of genes) in predicting annotated cell identity. Violins represent bootstrap distributions (n=100). **(c)**, Examples of spatial gene expression profiles of genes as quantified by HybISS. Examples of cell type markers and metabolic genes are showcased (upper and middle rows). Representative lipid distributions from MALDI-MSI data are shown (bottom row). **(d)**,. Cell clusters according to gene expression in different gene sets. Clusters were obtained independently for each gene set, and assigned to the annotated cell type with the most similar transcriptional profile. Only cells with at least 5 counts were considered for clustering, thus generating sparser clusters in the metabolic set. Lipid clusters were obtained independently and color-matched to anatomically similar transcriptional clusters. **(e )**, Bootstrap estimates for weighted average accuracy for classifiers using marker gene, metabolic gene, or lipids.

**Figure S15:**
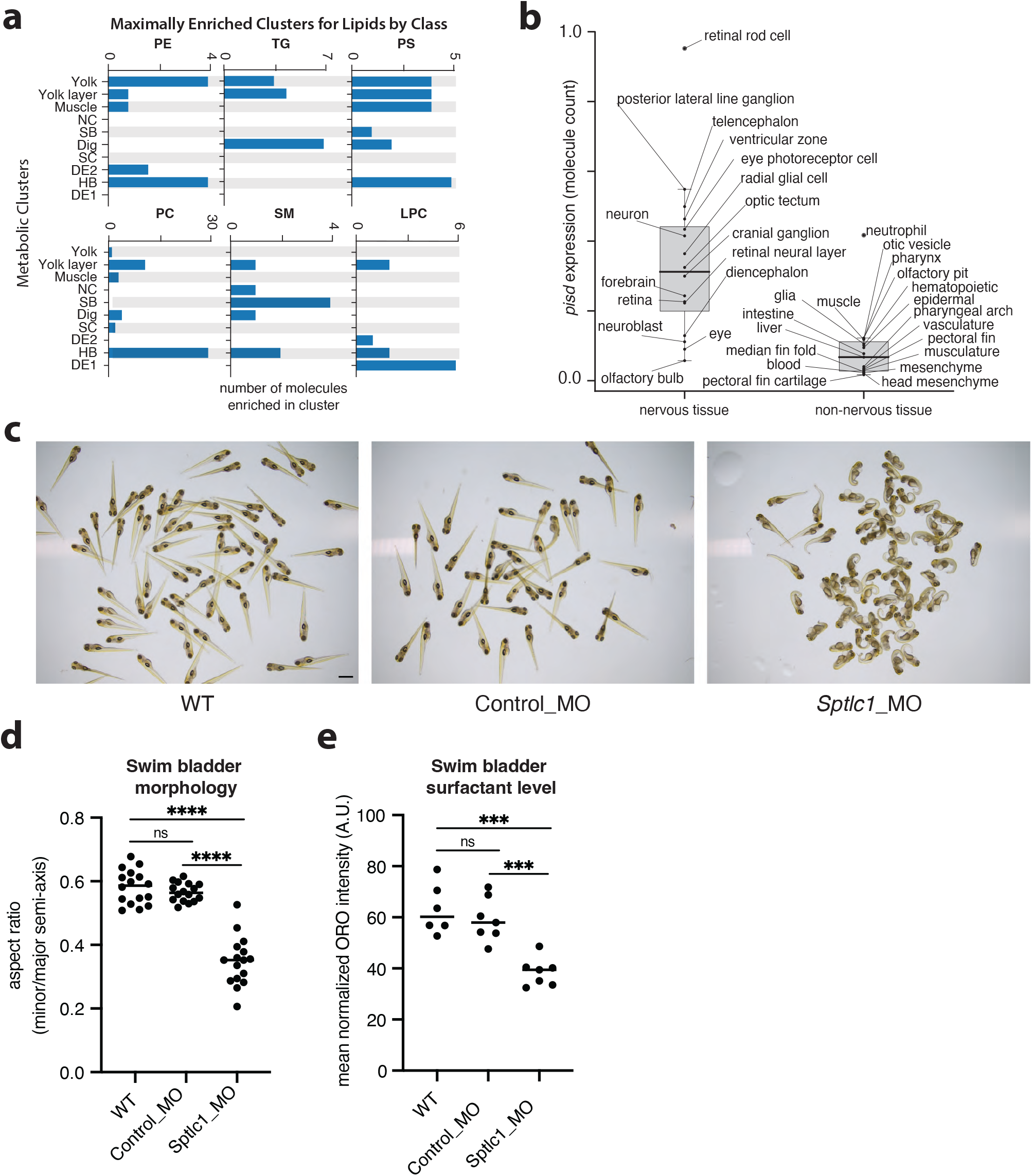
Metabolic organization of lipids in the 72 hpf zebrafish embryo. **(a)**, Barplots for each lipid class detailing the lipid enrichment of different clusters (i.e. metabolic region). Lipids are assigned to the metabolic region where they are maximally expressed. **(b)**, Barplot displaying the molecule counts of *Pisd* transcript in cell types defined from the scRNAseq dataset at 72 hpf zebrafish [48], data is stratified in nervous and non-nervous tissue. **(c)**, Representative images of 120 hpf zebrafish morphants for *Sptlc1* (Sptlc1-MO), control injected (Control-MO) and not-injected siblings (WT). Scale brar=1mm. **(d)**, Aspect ratio (minor/major semiaxis) of the swim bladder in larvae injected with control (Control_MO) or anti-Sptlc1 morpholinos (Sptlc1_MO) and not-injected siblings (WT); unpaired t-test, ****= p<0.0001(*Sptlc1*) zebrafish. **(e)**, Mean ORO intensity in the swim bladder in larvae injected with control (Control_MO) or anti-Sptlc1 morpholinos (Sptlc1_MO) and not-injected siblings (WT); unpaired t-test, ****= p<0.0001(*Sptlc1*) zebrafish.

## METHODS

### 1. Data generation

#### 1. Animal work and Cryosectioning

Zebrafish embryos of the Tubingen long fin (TL) strain were kept at 28 °C until collection at 8, 24, 48 and 72 hours post fertilization (*hpf* ). Following manual dechorionation, embryos were euthanized by fixation in 4% PFA at 4 °C overnight. After rinsing with 1X PBS, embryos were included in a block of 10% Porc-Skin gelatine (type A, Sigma), let solidify at 4 °C for 1h and then immediately frozen in isopentane pre-chilled to -65 °C. Specimens were kept at -80 °C until cryosectioning over the sagittal axis at -28 °C for a sectional thickness of 12 μm. Sections were stored at -80 °C until MALDI-MSI acquisition. All zebrafish husbandry procedures have been approved and accredited by the federal food safety and veterinary office of the canton of Vaud (VD-H23)

#### 2. MALDI-MSI datasets

Publicly available MALDI-MSI datasets were retrieved from Metaspace. Specifically, the human kidney samples were part of the NIH Kidney Precision Medicine Project (ID: 2024-06-06_19h26m13s, 2024-06-06_19h22m09s, 2024-06-06_19h16m21s, 2024-06-06_19h08m24s, 2024-06-06_19h03m38s, 2024-06-06_18h57m58s, 2024-06-06_18h45m39s, 2024-06-06_18h40m58s) and the baboon lung samples were in the Pacific Northwest National Laboratory (ID: 2024-06-11_18h30m14s, 2024-06-11_18h28m14s, 2024-06-11_18h35m41s, 2024-06-11_18h26m59s, 2024-06-11_18h43m25s). The datasets were chosen such that the acquisitions represented almost identical samples, allowing us to assume that the same molecules should be present at similar intensity levels. From each section in a given dataset, images corresponding to m/z values were retrieved on the Metaspace interface and intensity ranges were set equal for all sections. If an m/z was not identified by the interface, the image was marked as ‘missing’. Images with clear aberrations (i.e., missing pixels or significant differences in measured intensity between sections) were highlighted.

We collected MALDI-MSI datasets for zebrafish embryos. Sections from alternating sagittal sections were let dry at room temperature prior to coating with CHCA matrix prepared with 15 mg/μL in a solution acetone:water 1:1 and 0.1% trifluoroacetic acid (TFA) for positive ionisation. Matrix was deposited with a Sprayer (Sprangler) at 350 rpm and flow rate 5 μL/min for a total of 40 min. All MALDI-MSI acquisitions were acquired using an AP-SMALDI5 AF system coupled to a Q Exactive orbital trapping mass spectrometer in positive ion mode and in the 400-1200 m/z range. The resolution of acquisitions for the 24, 48 and 72 *hpf* embryos was 7 μm. For the 8 *hpf* embryo the resolution taken was 5 μm to compensate for its small size. In addition to these sections used for 3D image reconstruction, we also acquired sections from a biological replicate for each time point. The overall number of MS images acquired was 96.

#### 3. HybISS

Protocol was followed according to (Gyllborg et al., 2020) [47]. Zebrafish larvae were euthanized by fixation in 4% PFA at 4 °C overnight, embedded in 2% carboxymethyl cellulose (CMC) and cryosectioned at 10 μm. Sections were stored at −80 °C and processed within a week. For transcripts detection, sections were incubated with padlock probes (10-50 nM concentration per probe) for both marker and metabolic gene in the same mix. Samples were imaged on a Leica DMi8 epifluorescence microscope equipped with LED light source (Lumencor SPECTRA X, nIR, 90-10172), sCMOS camera (Leica DFC9000 GTC, 11547007) and 20x objective (HC PC APO, NA 0.8, air), with 10% overlap between tiles and 8-12 z-stack planes with 1 μm spacing.

Image processing: Z-projection was done using a variant of wavelet-based extended depth-of-field (EDF) (Forster et al., 2004). Tiles were stitched with ASHLAR (Muhlich et al., 2022) using the DAPI channel of the acquisition. Inter-cycle registrations were obtained using the wsireg Python package (Patterson et al. 2022), which wraps elastix (Klein et al., 2010). Spot detection was performed with Spotiflow (Dominguez Mantes et al., 2023) using the pretrained *HybISS* model in its default settings. Detections were linked (decoded) using a variant of Starfish’s (Axelrod et al., 2018) nearest neighbour decoder which uses the spot probabilities output by Spotiflow instead of the spot intensities.

#### 4. Bulk lipidomics experiments

Zebrafish embryos were collected at the desired developmental stage and anesthetized with addition of tricaine. After manual dechorionation, 15 individuals per replicate were pooled together and immediately frozen in liquid nitrogen. For each developmental time point (8, 24, 48 and 72 *hpf* ), 4 replicates were analyzed by high-throughput targeted HILIC MS/MS lipidomics workflow at facilities in the Université de Lausanne. Complex lipids were quantified using high-coverage, stable isotope dilution liquid chromatography-tandem mass spectrometry approach. Zebrafish tissue (15 embryos) was extracted with isopropanol (IPA) (Medina et al., 2020) pre-spiked with the internal standard (IS) mixture containing 75 isotopically labeled lipid species (with multiple representatives per lipid class, with varying fatty acid composition). The resulting extract was analyzed by hydrophilic interaction chromatography coupled to electrospray ionization tandem mass spectrometry approach (HILIC ESI-MS/MS) in positive and in negative ionization mode, using a TSQ Altis LC-MS/MS system (Thermo Scientific), as previously described by Medina et al. (2023) [45]. In two separate runs (using a dual-column setup), 1166 lipid species belonging to five major classes of complex lipids (glycerolipids, glycerophospholipids, cholesterol esters, sphingolipids and free fatty acids) were quantified with high precision and specificity. Using HILIC separation endogenous lipids co-elute with their corresponding IS thus allowing for the appropriate correction of matrix effects (in the same solvent composition) (Worlab et al., 2020; Lange et Fedorova, 2020). Optimized lipid class-dependent parameters were used for data acquisition in timed selected reaction monitoring (tSRM) mode. Raw LC-MS/MS data were processed using the Trace Finder Software (version 4.1, Thermo Fisher Scientific). The lipid abundance was reported as estimated concentration using a mixture of internal standards (IS) spiked at known concentrations and designed to correct for the differences in ionization and fragmentation efficiency depending on the fatty acid chain length and degree of unsaturation (Conterpois et al., 2018).

#### 5. Morpholino injection and Oil-Red-O staining

A translation-blocking morpholino (sptlc1-ATG MO: 5’(ACCCACTGTTGCCCCGACGCCATTT) [21]) (Gene Tools, Inc) was used to knock-down *Sptlc1* expression. Morpholinos were diluted at a concentration of 0.1 mM. A bolus of 100 um was injected in the cell of one-cell stage embryos of AB zebrafish strain. As a control, a mixture of random sequences was injected. Injected embryos were raised in an incubator at 28 °C with the addition of 1-phenyl 2-thiourea (PTU) to injibit melanine production to better visualize the swim bladder. The phenotype was assessed at 120 hpf. Non-injected siblings of the same clutch were also grown in the same conditions to control for any additional defects not related to the injections. At 120 hpf, about 8 representative larvae of each condition were placed in depression wells and photographed. Larvae were immediately fixed in 4 % PFA at 4 °C for 5 hours. Fixed zebrafish were then rinsed in PBS, incubated in MetOH at -20 °C for 24h, briefly equilibrated in 60% isopropanol and stained with a solution of 0.3% Oil-Red-O (Sigma-Aldrich (O0625))in 60% isopropanol for 3 hours as previously described ([20]. Samples were then rinsed with 60% isopropanol. After a final wash in PBS, organisms were placed in depression wells and photographed.

### 2. Models and Computational Approaches

The uMAIA framework comprises three main algorithms designed to achieve the following tasks:

- Peak calling for image generation from raw mass spectra
- Peak matching for the creation of a unified feature space
- Intensity normalization for batch effect removal

#### 1. Peak calling: processing of single MSI acquisitions

The first algorithm is concerned with processing the raw mass spectra within single MSI acquisitions in order to call peaks and construct images .

##### Peak calling of MSI spectra

Raw MSI data are typically stored as a collection of tuples of m/z values: its corresponding signal intensity and co-ordinates (*m*_*z*_, *intensity, x, y*). We, first, round m/z values to the nominal mass resolution of the instrument to obtain *N* values (bins of size 10e-5 were used). Then the data is loaded as a sparse matrix *S*_*p*,*m*_ containing the signal intensity values recorded at each pixel *p* ∈ [0, *P*] and each position in the mass spectrum indexes with *m* ∈ [0, *N*]. Next, a histogram representation of the data, **f**_*m*_, is obtained by counting the non-zero entries across the pixels, i.e. **f**_*m*_ = ∑_*p*_[*S*_*p*,*m*_ > 0]. A Gaussian kernel of small bandwidth (10e-4) is, then, applied to **f**_*m*_ to reduce the impact of noise. Then, the following 2-steps (Algorithms 1 and 2) adaptive peak-calling procedure is applied to retrieve the set of peaks *ℳ* and their boundaries *ℬ*.

##### Image retrieval and further processing

Images can be extracted from the raw data using the retrieved intervals. Specifically, we aggregate *S* according to the set of boundaries resulting from Algorithms 1 and 2. For each pixel *p* and each compound *c* we have 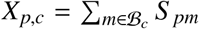. Images are total-ion content (TIC) normalized by dividing each pixel by the total signal detected: *X*_*pc*_ ← *X*_*pc*_/ ∑_*c*_ *X*_*pc*_.

Outputs of the peak-calling method are: (1) images saved in a .h5ad format where m/z values are variable names and (2) a .csv file specifying the intervals that were selected as well as the most frequent m/z value for the chosen bin among other metadata.

###### Algorithm 1 Retrieval of *ℳ*

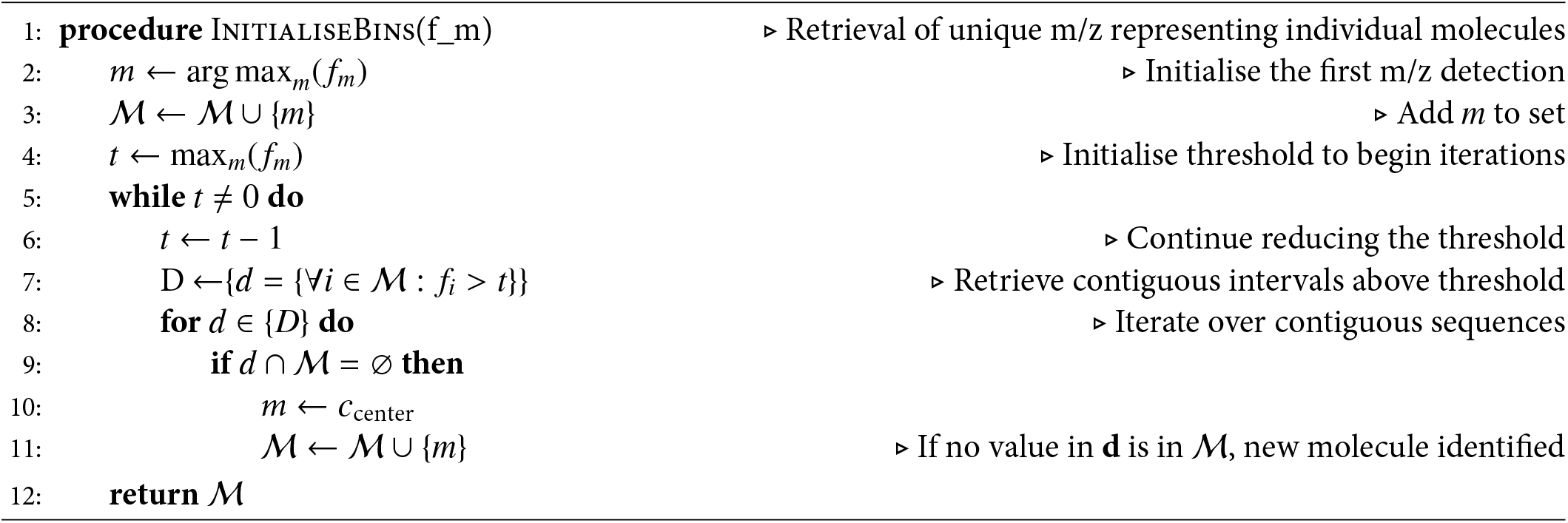

###### Algorithm 2 Determine the intervals *ℬ*

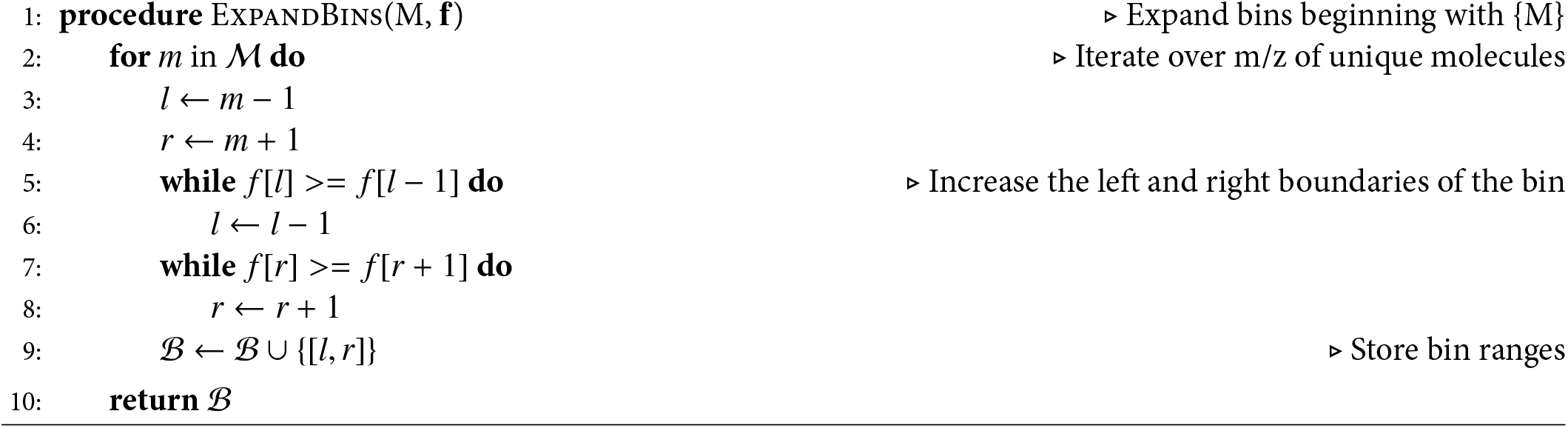

#### 2. Network flow-based peak matching: Optimized matching of peaks into a unified feature set

Mass spectra obtained from individual acquisitions differ in the number of identified peaks and their precise m/z position. Thus, MSI datasets are not directly produced in the same feature space. uMAIA approaches the creation of a unified feature space (featurization) in a way not contingent on an external reference by adopting a peak matching strategy: a peak from each acquisition is connected with one from another acquisition if they are considered the same molecule.

Connected peaks should satisfy a set of desired properties and some constraints. For example, closer peaks should be matched preferentially, one would rather match a peak than leave it unconnected, and each peak can be connected only to others that are also connected among themselves (as they are the same molecule). These properties are not trivially achievable with a heuristic or procedural approach such as building a nearest neighbor. Ideally, we aim to identify the choice of connections that respects all properties and constraints.

To formalize the problem, we consider the following quantities:

- Each of the *n* acquisitions provides a set of m/z peaks 𝒜 _1_, 𝒜 _2_, …, 𝒜 _*n*_. For simplicity, we refer to A as the union of all sets, with |𝒜 | = *L*.
- Considering each element in A as a node in a graph 𝒢, we define *G*_*ij*_, the adjacency matrix indicating the possible connections that are allowed. This is defined using a distance threshold *t* so that *G*_*i*, *j*_ = 1 if *dist*(*i, j*) < *t*, forcing *G*_*ij*_ = 0 if *i* and *j* are from the same acquisition.
- The desired matching uMAIA achieves is represented by a graph 𝒳 with an adjacency matrix *X*_*ij*_. *X*_*ij*_ = 1 when peaks from two acquisitions are considered to have originated from the same ion/compound. Each connected component of this graph is, then, considered a feature and used to construct the final feature space.
- *C*_*ij*_ indicates the associated cost of a particular matching *X*_*ij*_ = 1. The matrix is set by the distance in m/z between two peaks.

Overall, we set ourselves to find the optimal 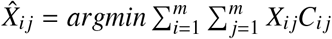 while also obeying a set of constraints. In its general form, the problem can be categorized as an Integer Linear Problem (ILP), which is only tractable for small *L* (while we have a large graph). However, two important simplifications we make here render it tractable in polynomial time. First, we divide the mass spectra into m/z intervals spanning 1 unit, which allows us to solve mz_range number of smaller subproblems (approximately 800). Second, we do not consider all the possible links *G*_*ij*_ between any acquisitions but allow only peaks between some pairs of acquisitions. Specifically, we considered the acquisitions ordered and allowed edges between the peaks of an acquisition and the *k* consecutive ones (for all the analyses k=2). Also, we introduce a start and end node in the graphs which can be connected with high cost *log*(*L*) but they are required by the constraint to complete a path. Since the solution of each subproblem can be found extremely fast using LP solvers, we solve it *M* times for *M* orders of the acquisitions obtained by random permutation and choose the best solution for each interval. The problem we solve can be formulated as follows:

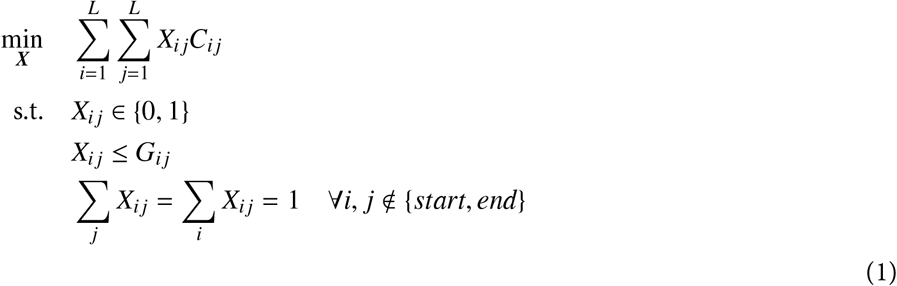

After solving the problem, features are retrieved by traversing all paths beginning from the start node until the end node is encountered.

#### 3. Normalization: intensity correction for MSI datasets

MSI experiments are prone to fluctuations of the detected signal intensity due to sample preparation and other experimental factors, potentially distorting the intensity histograms of different acquisitions. These technical intensity variations, or “batch effects”, can obscure real biological discoveries, reducing statistical power and complicating multivariate analyses.

Pixel intensity histograms from MSI data are typically bimodal, consisting of background and foreground signal distributions or components. Not all images contain both components. Our framework models batch effects in the foreground signals, assuming background signals are relatively stable.

Our approach to correct batch effects follows these steps:

- Estimation of the batch effect using a hierarchical Bayesian model.
- Use of the result of the estimation to determine signal intensity transformations for each image.
- Application of the functions to normalize the data.

##### Theory, model and implementation of the normalization

We consider the complete set of pixels *p* measured across all acquisitions *a* so that *p* ∈ ⋃ _*a*_ *𝒮*_*a*_, and denote *y*_*cp*_ the ground truth value associated to each compound *c* measured. Our goal is to correct batch effects and recover these ground truth values despite distortions *T*_*ca*_ induced by the measurement process for each acquisition *a* and compound *c* where we measure

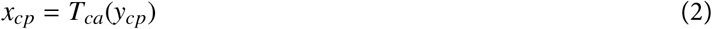

with *p* ∈ *𝒮*_*a*_. To compare images from different acquisitions, we aim to estimate 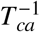 so that we can perform the transformation to correct the data.

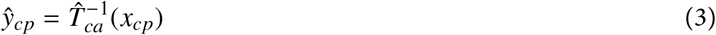

For this reason, we propose to estimate 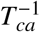 from the intensity value histograms observed across acquisition and compounds. We start by defining the “reference” intensity distribution across all pixels with a probability density function (PDF) *f*_*c*_(*y*_*cp*_) or equivalently, *f*_*c*_(*y*). The transformation *x*_*cp*_ = *T*_*ca*_(*y*_*cp*_) induces a new probability distribution *g*_*ca*_(*x*). The relation between *g, f* and *T* is particularly simple if we consider the cumulative density functions *G*_*ca*_ and *F*_*c*_ so that we can write

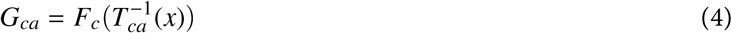

which immediately provides a way to estimate the desired normalizing transform 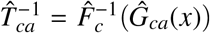. Since 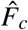 does not depend on the acquisition, it can be chosen with some freedom since all the normalization is performed by applying *Ĝ*_*ca*_ and F . We focus thus on the estimation of *Ĝ*_*ca*_, or equivalently, of *ĝ*_*ca*_.

Generally after parametrizing 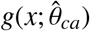, one would estimate the parameters 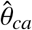 by maximum likelihood

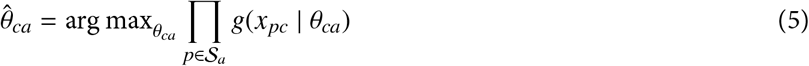

What complicates this endeavor is that the measured *x*_*pc*_ coming from a single acquisition cannot be expected to be representative of the complete set of pixels ⋃_*a*_ *𝒮*_*a*_. For example, in a real acquisition, there could be no (or very few) pixels with high signal while, in another, significantly more. In other words, if each acquisition were representative of all pixel intensity values across all acquisitions, the problem could be approached by histogram matching. However, since this is not the case, we need to combine information across acquisitions and compounds to obtain a robust 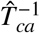.

To achieve this, we first consider the parameterization of *g* as a mixture of two Gaussians:

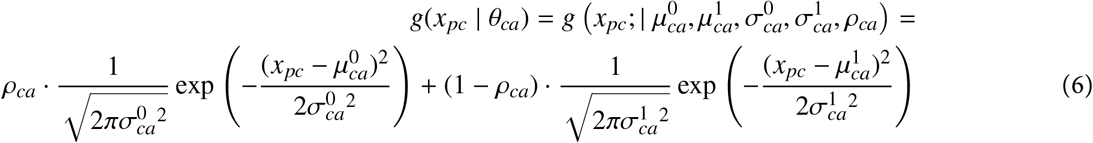

We assume that the parameters of the mixture can be factorized as to consider batch effect derived from acquisition- and compound-specific factors, given by *γ*_*a*_ and *λ*_*c*_, respectively. Specifically, we choose: 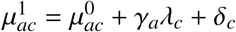 where *γ*_*a*_ ∈ [−2, 2] *λ*_*c*_ ∈ [−2, 2] and a prior for *δ*_*c*_ ∼ *Normal*(3, 1) represents the expected difference between modes. The prior for the background mode us 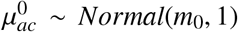 with *m*_0_ chosen in empirical Bayes fashion by considering the distribution of *σ*^1^ of a set of naive 2-component GMM fits. Similarly, 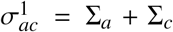. The following priors: Σ_*a*_, Σ_*c*_ ∼ *Exponential*(*s*) where *s* was chosen in empirical Bayes fashion by considering the distribution of *σ*^1^ of the full set of naive 2-component GMM fits.

Overall this corresponded to the following model:

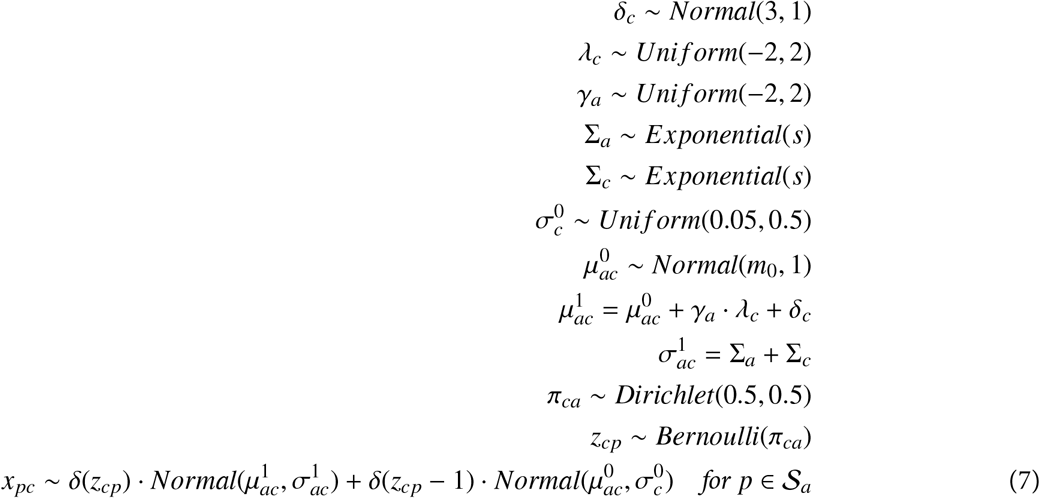

The model was implemented and fit using the *numpyro* probabilistic programming. The MAP estimates of the parameters 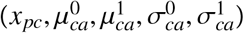 are determined and used to compute the transformation. *f* was chosen as the bi-modal Gaussian mixture model with the following parameters:

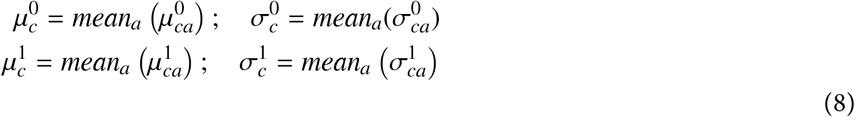

The CDF, *G*_*ca*_ and the inverse CDF 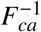 are evaluated numerically and used to compute the transformation 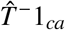.

### 3. Simulations and Method Evaluation

#### 1. Analysis of single acquisitions and Peakcalling

##### Comparison with other methods

A number of freely available software were applied and used for benchmarking. These included the following:

1. **MALDI-Quant**: available as an R package for MSI processing and calibration (based on adaptive binning) (Strimmer and Gibb, 2012)
2. **METASPACE**: online platform for visualisation and metabolite annotation (based on static binning and references m/z values) (Alexandrov et al., 2019)
3. **Mirion**: software for the visualisation of molecules in MSI datasets (based on static binning) (Paschke et al., 2013)
4. **MSI-Reader**: software for the visualisation of molecules in MSI datasets (based on static binning and references m/z values)

In order to image a molecule at spectra where it is detected, a range of m/z values that encompasses the mass shifts for the peak must be considered. Ranges identified by each method were retrieved and compared with the *uMAIA* method (Algorithms 1 and 2).

##### Evaluation of mass shifts within images

In order to estimate the degree to which mass shifts are correlated between molecules that are almost isobaric, m/z ranges were first manually established for each molecule in a single MALDI MSI acquisition and a reference m/z that corresponded to the most frequent m/z detection was identified. Next, for each molecule (i.e. m/z range), a mass shift for each spectrum was calculated by subtracting the observed m/z from the reference m/z. Mass shifts for different pairs of molecules were then scattered together for the subset of pixels that contained both molecules. Pearson’s R was calculated for the same subset. For visualisation, the vector was reshaped into its image dimensions.

##### Simulations for tested approaches

A simulation was setup to mimic mass shifts seen in real data. First, a set of numbers representing theoretical m/z values were generated in a unit range. To create datasets of variable molecular crowding, the size of the set of numbers was varied. To model mass shits, random values sampled from a Normal distribution with sigma=0.01 (as approximated from real data) were added to the theoretical m/z values for each of the 1000 simulated spectra in a way such that the order of the molecules after addition of noise still followed the original ordering of the molecules. In other words, two consecutive molecules cannot invert their positions in the range from one spectrum to another, as this would be inconsistent with what is possible in a real setting.

Next, noise ‘spike-ins’ were added at random across the range with variable intensities to reflect different Signal-to-Noise Ratios (SNR). An SNR of 10 in a spectrum with 100 detections would mean that there are 10 times as many real signals than noise, resulting in 90 true signals and 10 noise signals.

For each dataset that varied in its molecular crowding and SNR, 10 instances were simulated to better gauge the variability of the methods. Bins were retrieved by static binning methods (sizes of 0.02, 0.04), MALDI-Quant and uMAIA. Using the outputs, a dataframe was constructed to represent the target molecule, and the predicted molecule. Mutual information scores were calculated for each peak and a weighted average was reported.

##### Image quality assessment of MSI data

For each of the intervals returned by the different methods (uMAIA, Mirion, and MALDI-Quant), the fraction of generated images likely to contain multiple different molecules was estimated by counting images whose signal was contributed by more than a peak per spectrum/pixel.

For the analyses that required the intervals had to be aligned between different methods, we aligned them considering the extent of overlap between two ranges. With this information we calculated the following:

- the average intensity between two images referring to the same molecule from different methods
- the spatial correlation between the two images
- the number of uMAIA detections identified in individual MALDI-Quant ranges
- the number of MALDI-Quant detections identified in individual uMAIA ranges

We considered the number of MALDI-Quant detections that were within a given uMAIA range, and calculated the average mutual information for each molecule set.

To assess the quality of images retrieved by the different tested methods, 3 metrics were used to quantify various characteristics of image quality.

1. **Spatial chaos** considers intensities in an image and partitions them into levels. By assuming images arising from noise have equal distributions of intensity at each level, a score is assigned based on how different the sizes of the intensity partitions are. Codes for the spatial chaos metric were taken from https://github.com/alexandrovteam/pySM (Palmer et al., 2016)).
2. The **Noise model** metric was devised to distinguish images that have spatial structure. A basis function consisting of a set of small Gaussian-distributed points spanning the entire image was used, and a generalized linear model was fit to the image using a Poisson distribution to account for noise. A likelihood ratio test against a null hypothesis was used as a readout of the fit.
3. The **Power spectrum** metric was devised to quantify high-frequency aberrations across the image. Since MSI data are acquired by scanning, it is not uncommon to see aberrations that consist of stripes in the image. To quantify the presence of such high-frequency aberrations over both axes in the image, the image was converted into its Fourier space, and the area representing low and high frequency signals was summed, and their ratio was calculated.

A threshold was set for each metric based on manual inspection, and kept the same for all subsequent datasets that were analysed. These datasets included a DESI, FTICR, and 4 diverse samples from an APSMALDI instrument. The number of images that surpassed all thresholds for their respective metrics was calculated and shown for each dataset and method.

#### 2. Peak matching

##### Simulation to assess matching

To mimic mass shifts as seen in real data, the same simulation was done as described above. The number of sections was varied, and 10 realizations were taken. The ILP optimisation was run, and benchmarked to a binning approach where the size of the bin was equal to 2*std of specified noise distribution. Metrics including precision, accuracy and recall were calculated.

##### Evaluation of peak-matcher on real data

The peak calling algorithm was applied to 20 consecutive sagittal embryonic zebrafish sections. Theoretical m/z values were retrieved for all molecules in each section, which were then subjected to either our network flow matching method or traditional binning methods. Four different bin sizes (0.001, 0.0025, 0.005, 0.01) were used to group together compounds across the acquisitions.

The assessment of the detection accuracy was based on molecules we know should be present in the samples: matrix compounds and isotopologues. First, 50 of the most abundant matrix compounds across the acquisitions were considered. To assign an interval/peak to one of these compounds, we used the overlap with the theoretical m/z value as the criterion. For each individual acquisition, we considered a score of ambiguity computer by counting the number of observations falling inside the interval over the single one expected. Second, to score how consistently isotopologue pair (N, N+1) we present when featurizing with our approach vs standard binning, we considered the two binary vectors representing reporting the N and N+1 isotope presence respectively, and we computed both the Jaccard and Euclidean between the pair. This was performed only for the top 50 most abundant compounds to avoid stochastic dropouts occurring closer to the detection threshold, confounding the analysis.

To attain a more global perspective of the ability to featurize the data, we considered all molecules present across the acquisitions. 4 different bin sizes (0.001, 0.0025, 0.005, 0.01), including our ILP approach, were used to group together compounds across the sections.

#### 3. Batch effect characterization

##### Empirical estimation of the error matrix to study batch effects

To verify that the normalization model described above is justified by the error structure of the data (i.e., that fluctuations in the data are a combination of sample-dependent and feature-dependent factors), we consider a simplified empirical estimation framework to estimate the error. This empirical estimation is not generally applicable, as it exploits the atlas-sized data and the fact that our acquisitions are sequential sections of the same structure, but it was useful to explore the characteristics of the batch effect.

The estimation is based on these assumptions:

1. the observed background mode is equal to the true background mode 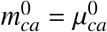.
2. As sections are consecutive, the high-frequency deviations in foreground mode shifts between consecutive sections are attributable to the batch effect, which we desire to remove. In other words, biological signals should vary smoothly across sections.

Following the following two equations from the model described above:

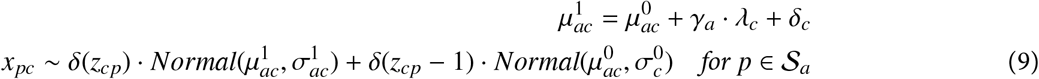

Assuming a simple factorization of the noise we can write a reparametrization:

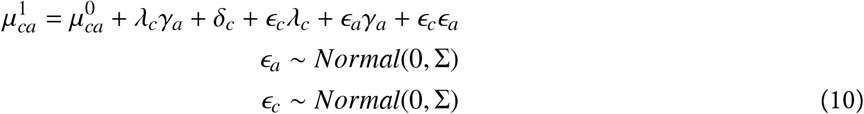

We can factor out from this expression the real biological signal *y*_*ca*_:

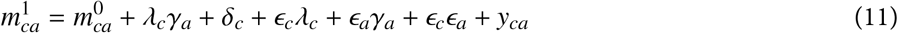

And considering this equation and the assumption above empirically estimate the variables 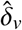and *ŷ*_*i*,*v*_

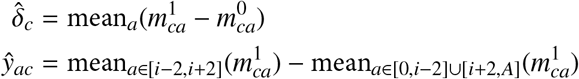

We can then approximate the matrix *M*_*ca*_ which quantifies the batch effect shift for each molecule and section.

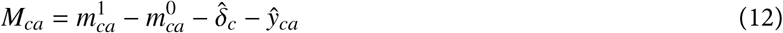

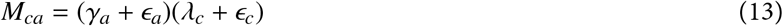

Singular Value Decomposition (SVD) was applied to the matrix to factorize it into two components representing acquisition- and molecule-dependent effects and the first eigenvalue from SVD is reported. Bootstrapped estimate of the first eigenvalue of this matrix was calculated to retrieve its variance by sampling 80% of molecules for 50 repetitions.

##### Simulation datasets for normalization method and evaluation

Two simulated datasets were created to assess the performance of various normalization algorithms. The first dataset evaluated the methods on a phantom object, while the second used real and complex distributions that were taken from ISH experiments of the mouse brain (Allen Brain Atlas).

The first dataset is an oversimplified tissue structure constructed as three ellipsoids in a 3D space, with two of the three ellipsoids partially overlapped. This simplicity facilitates the interpretation of the results. Multiple molecules were simulated to be present in one or multiple of the ellipsoids. When a molecule was assigned to an ellipsoid, pixel values in the region were drawn from a Normal distribution with a mean randomly generated from a Normal distribution. In cases where the molecule was present in ellipsoids that overlapped, the intensities were summed at the overlap, resulting in a third mode in the molecule’s intensity distribution. We simulated the sectioning of the data in a set of 2D images (sections).

Each section was then perturbed with batch effects to mimic the effect seen in the real data. Specifically, first factors representing section- and molecule-specific noise were drawn from a Normal distribution. Then, these parameters were used to derive and transform the intensities distributions from ground truth data to batch-effect affected (e.g., the inverse of the one we aim at estimating with our method).

A second simulation was performed using murine brain ISH images from the Allen Brain Atlas (ABA). Images were normalized by Min-Max rescaling to reflect the dynamic range present in MALDI-MSI data after logging the data ([-7,-1]). Next, batch effect noise was applied as was done above for the phantom object. Noise-perturbed data was then corrected using z-normalization, ComBat and uMAIA and quality of the correction was calculated via the Root Mean Square Error (RMSE) between corrected and ground truth data. Because treatment by the different normalization methods places the output data in different scales, a line was fit to the q-q plot between ground truth and corrected data. The slope of the line was used to place the two distributions in similar ranges so that intensity distributions within the image could be compared across the methods.

Using ground truth ABA ISH images and brain region annotations, we identified genes that revealed statistically significant differences between selected regions by application of two-sided t-tests (thalamus, hypothalamus, cortex, cerebellum and hippocampal formation). We compared these detections to significant results on batch-effect perturbed and normalized data by quantifying the number of true positive and false negative calls. Expected false positive and negative rates were estimated by bootstrapping (leaving 20% of the data)

##### Evaluation of algorithm on real data via clustering approach

We evaluated the unsupervised clustering of pixels before and after normalization and compared the result with a normalization algorithm (Combat) and imputation/integration algorithm (ComBat and scArches). All molecules that were present in at least 15% of the pixels were used as features for clustering. Clustering was performed with the K-means algorithm, and the number of clusters was set to 30.

##### Ability to perform regional differential intensity testing

ABA ISH data were used (refer to simulations in preceding section). For each gene, a differential expression test was made for all pairs of 5 selected regions by considering the set of pixels belonging to each region. For each tested pair, a matrix representing TP and TN values was constructed (TP identifed when p<0.05). The process was repeated for perturbed data, and data corrected using different methods (z-normalization, ComBat and uMAIA). FP, FN, TP and TN instances could then be ascertained when compared to ground truth. The ground truth False-Positive Rate and False-Negative Rate was estimated by bootstrapping sections belonging to the ground truth.

### 4. Zebrafish analyses

#### 1. Developmental analyses

##### Quantitative bulk LC/MS analysis

Data were converted to mol% (measured concentration / total lipid concentration). Averages across replicates for each lipid and time point were calculated. A UMAP of the lipids was computed, and annotated based on the time at which lipids were maximally abundant after dividing lipid concentration by their average so as to highlight relative changes in concentration between lipids.

##### Identification of spatially-informative molecules across timepoints

To identify spatially informative molecules, we performed ANOVA for each feature across all clusters. Specifically, images were clustered using K-mean using both annotated and un-annotated features. For each annotated lipid and cluster, we downsampled to 100 pixels and performed ANOVA across all clusters. Lipids were considered spatially informative when ANOVA returned a p-value below 0.05.

##### 3D metabolic atlas and multivariate analysis

Sagittal sections of zebrafish embryos at 8, 24, 48, and 72hpf were processed with uMAIA algorithms for image extraction, peak matching, and normalization. All molecules that had non-zero intensity values in at least 15% of pixels in 80% of sections were kept for downstream analysis.

Annotation of the detected peaks was achieved by considering the m/z values retrieved from bulk quantitative lipidomics experiments. We matched a MALDI compound to the annotation the nearest neighbor in m/z, considering only matches whose distance was below 0.008 Da.

To construct the volumetric data, consecutive slides were aligned using affine transforms. A 3D array was constructed, concatenating all the images, and a Gaussian filter was applied (sigma=0.4). The 3D array was then visualized in Napari.

Multivariate analyses and clustering are performed considering the following preprocessing. PCA was computed, and the first 10 components (explaining over 90% of the dataset’s variance) were considered. Clustering was performed using the K-Means clustering algorithm to retrieve groups of voxels that were then characterized by lipid content and compared to known anatomical structures.

##### Computationally tracing metabolic pseudolineages

To link metabolic territories together across the developmental time points, we constructed a custom pairwise squared-Euclidean distance matrix between clusters considering the sequence of time points. Specifically, we start by initializing the distance matrix, *D*_*ij*_ to a large value (2.5). Next, we consider consecutive time points t and t+1, and we subset the lipids that were detected at both time points so to obtain two matrices of cluster averages 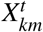 and 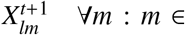 feat_*t*_ *& m* ∈ feat_*t*+1_ and squared-Euclidean distances between these pairs were computed and set as values of *D*_*ij*_ so that 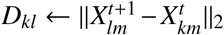. Then the *D*_*ij*_ was sorted by optimal leaf ordering. We show clusters in that order and display them to be spaced proportionally to the distance between consecutive clusters. In order to link together clusters from different time points. The Hungarian Algorithm was applied to match the clusters of each pair of timepoints *t* and *t* + 1. To identify the lipids that may play more important roles in branching or splitting of the lineages, we trained a decision tree classifier (scikit-learn implementation) to classify pixels belonging to one of 2 categories after branching. The feature importance scores we reported are bootstrapping averages.

#### 2. 3D metabolic atlas and multivariate analysis

20 sagittal sections corresponding to the zebrafish at 72 hpf were provided to the uMAIA framework for image extraction, peak matching and normalization. Prior to peak matching, the lowest 2 percentile of intensity peaks were removed. After normalization, all molecules that had non-zero intensity values in at least 15% of pixels in 80% of sections were kept for downstream analysis. Annotation of the detected peaks was achieved by consulting the m/z values retrieved from bulk quantitative lipidomics experiments, which were matched to our dataset by considering the closest match in m/z, within a threshold of 0.008Da. Individual images corresponding to lipids were rescaled to their correct dimensions and manually adjusted using linear transforms, i.e. side-to-side shifts and rotations to approximately align all the sections. A 3D array was constructed, over which a minor Gaussian blur was added (sigma=0.4). An outline of the fish defined by thresholding the sum of intensities over all molecules was made to show the volumetric bounds of the animal. The 3D array was then visualised in Napari with a custom heatmap.

##### Lipid-lipid correlation, enrichment and graph representation

A pairwise correlation distance matrix was calculated across lipids and visualized after column-row sorting using the SPIN algorithm (Tsafrir et al., 2005).

To construct the cluster-enrichment heatmap, we applied K-Means and, for each cluster, averaged the intensity for each lipid, resulting in a matrix with rows (dimension equal to the number of regions) and columns (dimension equal to the number of lipids). Each column was min-max normalised in order to highlight the region in which specific lipids are most abundant in.

To obtain the graph representation of the lipid-lipid similarity at each timepoint, we calculated a cluster-enrichment score for each lipid species. Pairwise Euclidean distances between enrichment scores distributions were computed. The matrix of distances was used to compute a force-directed layout. To improve the visualization, the 500 edges with the smallest distances were considered.

#### 3. Selection of metabolic genes, scRNAseq and HybISS analysis

To identify zebrafish genes involved in metabolic processes, we utilized the KEGG database, including genes associated with enzymatic reactions and metabolite transport (Kanehisaa and Goto, 2000). Single-cell transcriptomics data of zebrafish embryos at 72 hpf were obtained from the Zebrahub database (Lange et al., 2023). We selected three subsets of genes: the 50 most variable genes, the 50 most variable metabolic genes, and 50 random genes. For each subset, a training set of total cells was used to fit a linear discriminant analysis (LDA) model to predict cell types. Predictive accuracy was assessed over 100 bootstrap iterations.

HybISS data analysis: For each section, individual transcripts were binned into 13.6 μm (40 px^2^) areas, serving as proxies for cells. Genes with fewer than 5 counts and meta-cells with fewer than 5 genes were excluded. We used the Scanpy Python package (Wolf et al., 2018) to cluster cells with similar transcriptional profiles and Squidpy (Palla et al., 2022) to generate spatial plots of these clusters. Cluster annotation was performed by correlating the mean expression of HybISS cluster with that of annotated clusters from the scRNAseq dataset, selecting clusters with the highest correlation scores. The analysis was repeated using only marker genes or metabolic genes.

To compare the information content between marker genes, metabolic genes, and lipids, HybISS data were aligned with lipid data using Fiji. We considered all marker gene expression distributions, performed PCA across the feature space, and applied Leiden clustering to the top 10 principal components (Traag et al., 2019). Next, we retrieved bootstrap (n=35) estimates of each set’s (marker genes, metabolic genes, lipid distributions) ability to predict cluster identity: decision trees (scikit-learn implementation) with shallow depths (4) were used as classifiers.

#### *uMAIA* Package

The *uMAIA* is coded in python and its source is made available at https://github.com/lamanno-epfl/uMAIA.

## References

[1] D. Bressan, G. Battistoni, and G. J. Hannon. “The dawn of spatial omics”. en. In: Science 381.6657 (2023), eabq4964.

[2] S. R. Srivatsan et al. “Embryo-scale, single-cell spatial transcriptomics”. en. In: Science 373.6550 (2021), pp. 111–117.

[3] Y. Liu et al. “High-Spatial-Resolution Multi-Omics Sequencing via Deterministic Barcoding in Tissue”. In: Cell 183.6 (2020), 1665–1681.e18.

[4] C. Ortiz et al. “Molecular atlas of the adult mouse brain”. en. In: Sci Adv 6.26 (2020), eabb3446.

[5] A. Chen et al. “Spatiotemporal transcriptomic atlas of mouse organogenesis using DNA nanoball-patterned arrays”. en. In: Cell 185.10 (2022), 1777–1792.e21.

[6] Z. Yao et al. “A high-resolution transcriptomic and spatial atlas of cell types in the whole mouse brain”. en. In: Nature 624.7991 (2023), pp. 317–332.

[7] S. E. Flaherty et al. “A lipase-independent pathway of lipid release and immune modulation by adipocytes”. In: Science 363.6430 (2019), pp. 989–993.

[8] J. L. Anderson, J. D. Carten, and S. A. Farber. “Zebrafish lipid metabolism: from mediating early patterning to the metabolism of dietary fat and cholesterol”. en. In: Methods Cell Biol. 101 (2011), pp. 111–141.

[9] E. M. Storck, C. Özbalci, and U. S. Eggert. “Lipid Cell Biology: A Focus on Lipids in Cell Division”. en. In: Annu. Rev. Biochem. 87 (2018), pp. 839–869.

[10] E. R. Moellering, B. Muthan, and C. Benning. “Freezing tolerance in plants requires lipid remodeling at the outer chloroplast membrane”. en. In: Science 330.6001 (2010), pp. 226–228.

[11] J. C. M. Holthuis and A. K. Menon. “Lipid landscapes and pipelines in membrane homeostasis”. en. In: Nature 510.7503 (2014), pp. 48–57.

[12] A. L. Santos and G. Preta. “Lipids in the cell: organisation regulates function”. en. In: Cell. Mol. Life Sci. 75.11 (2018), pp. 1909–1927.

[13] S. A. Lim et al. “Lipid metabolism in T cell signaling and function”. en. In: Nat. Chem. Biol. 18.5 (2022), pp. 470–481.

[14] C. K. Glass and J. M. Olefsky. “Inflammation and Lipid Signaling in the Etiology of Insulin Resistance”. In: Cell Metab. 15.5 (2012), pp. 635–645.

[15] L. Capolupo et al. “Sphingolipids control dermal fibroblast heterogeneity”. en. In: Science 376.6590 (2022), eabh1623.

[16] X. L. Guan et al. “Biochemical membrane lipidomics during Drosophila development”. en. In: Dev. Cell 24.1 (2013), pp. 98–111.

[17] L. Zhang et al. “Low-input lipidomics reveals lipid metabolism remodelling during early mammalian embryo development”. en. In: Nat. Cell Biol. (2024).

[18] X. Zhu et al. “Advances in MALDI Mass Spectrometry Imaging Single Cell and Tissues”. en. In: Front Chem 9 (2021), p. 782432.

[19] M. Aichler and A. Walch. “MALDI Imaging mass spectrometry: current frontiers and perspectives in pathology research and practice”. en. In: Lab. Invest. 95.4 (2015), pp. 422–431.

[20] D. Fraher et al. “Zebrafish Embryonic Lipidomic Analysis Reveals that the Yolk Cell Is Metabolically Active in Processing Lipid”. In: Cell Rep. 14.6 (2016), pp. 1317–1329.

[21] I. Castanon et al. “Wnt-controlled sphingolipids modulate Anthrax Toxin Receptor palmitoylation to regulate oriented mitosis in zebrafish”. en. In: Nat. Commun. 11.1 (2020), pp. 1–14.

[22] X. Zhao et al. “Lipid alterations during zebrafish embryogenesis revealed by dynamic mass spectrometry profiling with C=C specificity”. en. In: J. Am. Soc. Mass Spectrom. 30.12 (2019), pp. 2646–2654.

[23] V. Pirro et al. “Lipid dynamics in zebrafish embryonic development observed by DESI-MS imaging and nanoelectrospray-MS”. In: Mol. Biosyst. 12.7 (2016), pp. 2069–2079.

[24] M. E. Dueñas, J. J. Essner, and Y. J. Lee. “3D MALDI mass spectrometry imaging of a single cell: Spatial mapping of lipids in the embryonic development of zebrafish”. en. In: Sci. Rep. 7.1 (2017), p. 14946.

[25] G. D’Angelo and G. La Manno. “The lipotype hypothesis”. en. In: Nat. Rev. Mol. Cell Biol. 24.1 (2023), pp. 1–2.

[26] B. Balluff et al. “Batch Effects in MALDI Mass Spectrometry Imaging”. en. In: J. Am. Soc. Mass Spectrom. 32.3 (2021), pp. 628–635.

[27] M. Tuck et al. “MALDI-MSI Towards Multimodal Imaging: Challenges and Perspectives”. en. In: Front Chem 10 (2022), p. 904688.

[28] D. Unsihuay, D. Mesa Sanchez, and J. Laskin. “Quantitative Mass Spectrometry Imaging of Biological Systems”. en. In: Annu. Rev. Phys. Chem. 72 (2021), pp. 307–329.

[29] A. McCann et al. “Mass shift in mass spectrometry imaging: comprehensive analysis and practical corrective workflow”. In: Anal. Bioanal. Chem. 413.10 (2021), pp. 2831–2844.

[30] J. O. Eriksson et al. “MSIWarp: A General Approach to Mass Alignment in Mass Spectrometry Imaging”. en. In: Anal. Chem. 92.24 (2020), pp. 16138–16148.

[31] J. Romson and Å. Emmer. “Mass calibration options for accurate electrospray ionization mass spectrometry”. In: Int. J. Mass Spectrom. 467 (2021), p. 116619.

[32] A. R. Buchberger et al. “Mass Spectrometry Imaging: A Review of Emerging Advancements and Future Insights”. In: Anal. Chem. 90.1 (2018), pp. 240–265.

[33] M. Pérez-Cova et al. “MSroi: A pre-processing tool for mass spectrometry-based studies”. In: Chemometrics Intellig. Lab. Syst. 215 (2021), p. 104333.

[34] C. Paschke et al. “Mirion—A Software Package for Automatic Processing of Mass Spectrometric Images”. In: J. Am. Soc. Mass Spectrom. 24.8 (2013), pp. 1296–1306.

[35] S. Gibb and K. Strimmer. “MALDIquant: a versatile R package for the analysis of mass spectrometry data”. en. In: Bioinformatics 28.17 (2012), pp. 2270–2271.

[36] A. Palmer et al. “FDR-controlled metabolite annotation for high-resolution imaging mass spectrometry”. In: Nat. Methods 14.1 (2017), pp. 57–60.

[37] M. Sprang, M. A. Andrade-Navarro, and J.-F. Fontaine. “Batch effect detection and correction in RNA-seq data using machine-learning-based automated assessment of quality”. en. In: BMC Bioinformatics 23.Suppl 6 (2022), p. 279.

[38] Y. Zhang, G. Parmigiani, and W. E. Johnson. “ComBat-seq: batch effect adjustment for RNA-seq count data”. en. In: NAR Genom. Bioinform. 2.3 (2020), lqaa078.

[39] W. E. Johnson, C. Li, and A. Rabinovic. “Adjusting batch effects in microarray expression data using empirical Bayes methods”. en. In: Biostatistics 8.1 (2006), pp. 118–127.

[40] X. Yu et al. “Batch alignment of single-cell transcriptomics data using deep metric learning”. en. In: Nat. Commun. 14.1 (2023), p. 960.

[41] Y. Zhang and F. Wang. “SSBER: removing batch effect fo single-cell RNA sequencing data”. en. In: BMC Bioinformatics 22.1 (2021), p. 249.

[42] M. Andreatta and S. J. Carmona. “STACAS: Sub-Type Anchor Correction for Alignment in Seurat to integrate single-cell RNA-seq data”. en. In: Bioinformatics 37.6 (2021), pp. 882–884.

[43] E. S. Lein et al. “Genome-wide atlas of gene expression in the adult mouse brain”. en. In: Nature 445.7124 (2006), pp. 168–176.

[44] M. Lotfollahi et al. “Mapping single-cell data to reference atlases by transfer learning”. en. In: Nat. Biotechnol. 40.1 (2021), pp. 121–130.

[45] J. Medina et al. “Omic-Scale High-Throughput Quantitative LC–MS/MS Approach for Circulatory Lipid Phenotyping in Clinical Research”. In: Anal. Chem. 95.6 (2023) pp. 3168–3179.

[46] C. B. Kimmel et al. “Stages of embryonic development of the zebrafish”. en. In: Dev. Dyn. 203.3 (1995), pp. 253–310.

[47] D. Gyllborg et al. “Hybridization-based In Situ Sequencing (HybISS): spatial transcriptomic detection in human and mouse brain tissue”. 2020.

[48] M. Lange et al. “Zebrahub – Multimodal Zebrafish Developmental Atlas Reveals the State-Transition Dynamics of Late-Vertebrate Pluripotent Axial Progenitors”. en. 2023.

[49] M. Kannan et al. “Phosphatidylserine synthesis at membrane contact sites promotes its transport out of the ER”. en. In: J. Lipid Res. 58.3 (2017), pp. 553–562.

[50] D. Giovannone et al. “Programmed conversion of hypertrophic chondrocytes into osteoblasts and marrow adipocytes within zebrafish bones”. en. In: Elife 8 (2019).

[51] W. Zheng et al. “Comparative transcriptome analyses indicate molecular homology of zebrafish swimbladder and mammalian lung”. en. In: PLoS One 6.8 (2011), e24019.

[52] C. B. Daniels et al. “The origin and evolution of the surfactant system in fish: insights into the evolution of lungs and swim bladders”. en. In: Physiol. Biochem. Zool. 77.5 (2004), pp. 732–749.

[53] T. Chen et al. “Development of the Swimbladder Surfactant System and Biogenesis of Lysosome-Related Organelles Is Regulated by BLOS1 in Zebrafish”. en. In: Genetics 208.3 (2018), pp. 1131–1146.

[54] C. L. Winata et al. “Development of zebrafish swimbladder: The requirement of Hedgehog signaling in specification and organization of the three tissue layers”. In: Dev. Biol. 331.2 (2009), pp. 222–236.

[55] Y. A. Hannun and L. M. Obeid. “Sphingolipids and their metabolism in physiology and disease”. en. In: Nat. Rev. Mol. Cell Biol. 19.3 (2018), pp. 175–191.

[56] A. L. Lillie RD. “Supersaturated solutions of fat stains in dilute isopropanol for demonstration of acute fatty degeneration not shown by Herxheimer’s technique”. In: Arch Pathol 36.1 (1943), pp. 432–440.

[57] I. Levental and E. Lyman. “Regulation of membrane protein structure and function by their lipid nano-environment”. en. In: Nat. Rev. Mol. Cell Biol. 24.2 (2023), pp. 107–122.

[58] R. Covino et al. “A eukaryotic sensor for membrane lipid saturation”. en. In: Mol. Cell 63.1 (2016), pp. 49–59.

[59] P. A. Janmey and P. K. J. Kinnunen. “Biophysical properties of lipids and dynamic membranes”. en. In: Trends Cell Biol. 16.10 (2006), pp. 538–546.

